# Integrated spatial transcriptomics and lipidomics of precursor lesions of pancreatic cancer identifies enrichment of long chain sulfatide biosynthesis as an early metabolic alteration

**DOI:** 10.1101/2023.08.14.553002

**Authors:** Marta Sans, Yihui Chen, Fredrik I. Thege, Rongzhang Dou, Jimin Min, Michele Yip-Schneider, Jianjun Zhang, Ranran Wu, Ehsan Irajizad, Yuki Makino, Kimal I. Rajapakshe, Mark W. Hurd, Ricardo A. León-Letelier, Jody Vykoukal, Jennifer B. Dennison, Kim-Anh Do, Robert A. Wolff, Paola A. Guerrero, Michael P. Kim, C Max Schmidt, Anirban Maitra, Samir Hanash, Johannes F. Fahrmann

## Abstract

**Background:** The development of diverse spatial profiling technologies has provided an unprecedented insight into molecular mechanisms driving cancer pathogenesis. Here, we conducted the first integrated cross-species assessment of spatial transcriptomics and spatial metabolomics alterations associated with progression of intraductal papillary mucinous neoplasms (IPMN), *bona fide* cystic precursors of pancreatic ductal adenocarcinoma (PDAC).

**Methods:** Matrix Assisted Laster Desorption/Ionization (MALDI) mass spectrometry (MS)-based spatial imaging and Visium spatial transcriptomics (ST) (10X Genomics) was performed on human resected IPMN tissues (N= 23) as well as pancreata from a mutant *Kras;Gnas* mouse model of IPMN. Findings were further compared with lipidomic analyses of cystic fluid from 89 patients with histologically confirmed IPMNs, as well as single-cell and bulk transcriptomic data of PDAC and normal tissues.

**Results:** MALDI-MS analyses of IPMN tissues revealed long-chain hydroxylated sulfatides, particularly the C24:0(OH) and C24:1(OH) species, to be selectively enriched in the IPMN and PDAC neoplastic epithelium. Integrated ST analyses confirmed that the cognate transcripts engaged in sulfatide biosynthesis, including *UGT8, Gal3St1*, and *FA2H*, were co-localized with areas of sulfatide enrichment. Lipidomic analyses of cystic fluid identified several sulfatide species, including the C24:0(OH) and C24:1(OH) species, to be significantly elevated in patients with IPMN/PDAC compared to those with low-grade IPMN. Targeting of sulfatide metabolism via the selective galactosylceramide synthase inhibitor, UGT8-IN-1, resulted in ceramide-induced lethal mitophagy and subsequent cancer cell death *in vitro*, and attenuated tumor growth of mutant *Kras;Gnas* allografts. Transcript levels of *UGT8* and *FA2H* were also selectively enriched in PDAC transcriptomic datasets compared to non-cancerous areas, and elevated tumoral *UGT8* was prognostic for poor overall survival.

**Conclusion:** Enhanced sulfatide metabolism is an early metabolic alteration in cystic pre-cancerous lesions of the pancreas that persists through invasive neoplasia. Targeting sulfatide biosynthesis might represent an actionable vulnerability for cancer interception.

## Introduction

Pancreatic cysts occur in up to 13% of patients undergoing abdominal imaging (CT scan or MRI) for reasons unrelated to pancreatic symptoms^1,2^. With an increasing use of abdominal imaging, the prevalence of incidental detection of pancreatic cysts continues to rise, with almost ∼500,000 new pancreatic cysts diagnosed each year in the United States alone. Retrospective histopathology studies on surgically resected pancreatic cysts have shown that approximately half are neoplastic entities. The most common cystic neoplasm that is a *bona fide* precursor to pancreatic ductal adenocarcinoma (PDAC) is intraductal papillary mucinous neoplasm (IPMN). IPMN is a mucin-secreting neoplasm arising in either the main pancreatic duct or one of the branch ducts and comprise roughly 40-50% of resected lesions that are initially diagnosed as asymptomatic pancreatic cysts^3,4^. IPMNs are lined by either low-grade (LG) or high-grade (HG) epithelial dysplasia, and in a subset of cases, histological progression culminates in cancer. Once an IPMN develops an invasive component, the probability of long-term survival drops substantially. Evolutionary modeling studies have suggested an average window of over three years between a pre-malignant IPMN and the development of HG dysplasia and PDAC^5^, providing an opportunity for cancer interception efforts^6,7^. Currently surgical resection is the only available definitive treatment modality for IPMNs, highlighting a pressing need to identify molecular vulnerabilities that would substantially delay or eliminate the progression of these pancreatic early lesions to invasive PDAC^8^.

Metabolic reprogramming is inherently intertwined with oncogenic drivers that promote malignant progression of pre-cancerous lesions of the pancreas^9–15^. For instance, oncogenic KRas in pancreatic acinar cells increases ATP-citrate lyase (ACLY)-dependent acetyl coenzyme A (CoA) production that is used in the mevalonate pathway to support acinar-to-ductal metaplasia, a precursor to PDAC^10,11^. Mutant *GNAS*^R201C^ cooperates with *KRAS*^G12D^ and *TP53* mutations to drive tumor initiation and progression to malignancy through protein kinase A (PKA)-mediated suppression of salt-inducible kinases (SIK1-3) that is accompanied by profound lipid remodeling and induction of fatty acid oxidation^12^. Elucidation of specific metabolic adaptations in pre-cancerous lesions that enable or accompany malignancy has potential to lead to the identification of actionable metabolic vulnerabilities with preventative or therapeutic potential.

The emergence of spatial ‘omics’ technologies have enabled unprecedented advances in our understanding of the molecular mechanisms that drive cancer pathogenesis^16^. In the current study, we performed an integrated cross-species assessment of spatial metabolomics and spatial transcriptomics alterations associated with progression of IPMN. Specifically, Matrix Assisted Laster Desorption/Ionization (MALDI) mass spectrometry-based imaging was performed on 23 surgically resected human IPMN and IPMN-associated PDAC cases, which led to the discovery of distinct enrichment of long-chain hydroxylated sulfatides in neoplastic epithelium. Leveraging our recent spatial transcriptomic (ST) dataset in IPMNs obtained via the 10x Genomics Visium platform^17^ for serial sections derived from the same tissues evaluated via MALDI-MS imaging, we demonstrated elevated transcripts encoding for enzymes involved in sulfatide biosynthesis to co-localize with enrichment of long-chain hydroxylated sulfatides found in epithelium lining the IPMN. These findings were validated in a genetically engineered mouse model of IPMN, co-expressing mutant *Kras;Gnas*^18^. Lipidomic analyses of cystic fluid from patients with IPMN further demonstrated elevated sulfatides to be associated with malignant progression. Targeting of sulfatide metabolism using a selective small molecule inhibitor of galactosylceramide synthase (CGT, also known as UGT8) decreased cell viability of *Kras;Gnas* cell lines and attenuated tumor development in a allograft model of IPMN. Interrogation of published single-cell RNAseq and bulk transcriptomic datasets for PDAC and normal pancreas tissues also demonstrated *UGT8* and *FA2H* transcript levels to be selectively elevated in PDAC. Elevated tumoral *UGT8* mRNA expression was found to be prognostic for poor overall survival.

## MATERIALS AND METHODS

### Tissue Specimens

Patient and murine sample collection for MALDI analyses Archival snap frozen (**Table S1**) tissue samples from 34 IPMN resections at either MDACC or Indiana University School of Medicine were sectioned, stained with hematoxylin & eosin (H&E) and evaluated by a pancreas pathologist (A.M) to identify areas containing IPMN lesions and cancer. The analyzed samples were comprised of 15 LG IPMN (all gastric), 4 HG IPMN, and 4 PDAC arising in the backdrop of IPMN. Tissues from 9 additional PDAC cases without a concurrent IPMN were also analyzed. Samples were cryo-sectioned onto Visium Spatial Gene Expression slides for ST analysis, as previously published^19^. For the murine samples, pancreata were collected from p48-Cre; LSL-KrasG12D; Rosa26RLSL-rtTA-TetO-GnasR201C mice (*Kras;Gnas* mice) at 25 weeks on doxycycline diet. Samples were flash frozen in liquid nitrogen, cryo-sectioned and H&E stained.

### Cystic fluid

Samples were obtained from the Indiana University Pancreatic Tissue-Fluid Bank following approval by Indiana University Institutional Review Board. Patients signed informed consent for collection of pancreatic fluid at the time of routine endoscopy (EUS or ERCP) and/or resection. Fluid specimens were placed immediately on ice after procurement and aliquoted for storage at −80 degrees. All samples were histopathologically confirmed as IPMN following surgical resection, and dysplastic grade was determined according to the World Health Organization (WHO) criteria.

### MALDI-MS imaging

Matrix-assisted laser desorption/ionization mass spectrometry (MALDI-MS) imaging was conducted on 38 pancreatic tissue sections from 34 patients. Tissue samples were cryosectioned at 10µm and stored at −80°C before use. Prior to analysis, tissue samples were sprayed with a solution of 10 mg/mL of 9-Amminoacridine in 70% methanol using the HTX M5 Sprayer (HTX imaging). The spray nozzle was heated to 75°C and the slide to 30°C. MALDI-MS imaging was conducted on a MALDI Synapt G2-Si (Waters, USA) at 60 µm spatial resolution. Data was collected in resolution and negative mode, from *m/z* 50-2000. After MALDI-MS imaging, the same and an adjacent tissue section were stained with hematoxylin and eosin (H&E) for pathological annotation. MALDI-MS imaging was performed on serial sections from a subset of the same tissue samples used for spatial transcriptomics using the Visium technology in our recently study^17^, thereby allowing us to directly compare the localization of the RNA transcripts and the sulfatides observed on the same tissue sample.

### Metabolomic analysis

#### Sample Extraction

Pre-aliquoted cystic fluid samples (10µL) were extracted with 30µL of LCMS grade 2-propanol (ThermoFisher) in a 96-well microplate (Eppendorf). Plates were heat sealed, vortexed for 5min at 750 rpm, and centrifuged at 2000 x g for 10 minutes at room temperature. The supernatant (10µL) was carefully transferred to a 96-well plate, leaving behind the precipitated protein. The supernatant was further diluted with 90µL of 1:3:2 100mM ammonium formate, pH3 (Fischer Scientific): LCMS grade acetonitrile (ThermoFisher): LCMS grade 2-propanol (ThermoFisher) and transferred to a 384-well microplate (Eppendorf) for lipids analysis using LCMS.

Cell lysates were washed 2x with pre-chilled 0.9% NaCl followed by addition of 2.5mL of pre-chilled 3:1 isopropanol:ultrapure water. Cells were scraped using a 25cm Cell Scraper (Sarstedt) in extraction solvent and transferred to a 15mL conical tube (Eppendorf). Samples were briefly vortexed followed by centrifugation at 4°C for 10 min at 2,000 x g. Thereafter, 1.2mL of metabolite extracts were transferred to 1.5mL Eppendorf tubes and stored in −20°C until metabolomic analysis.

#### Untargeted Analysis of Complex Lipids

Untargeted lipidomic analyses were conducted on a Waters Acquity™ UPLC system coupled to a Xevo G2-XS quadrupole time-of-flight (qTOF) mass spectrometer. Chromatographic separation was performed using a C18 (Acquity™ UPLC HSS T3, 100 Å, 1.8 µm, 2.1×100mm, Water Corporation, Milford, U.S.A) column at 55°C. The mobile phases were (A) water, (B) acetonitrile, (C) 2-propanol and (D) 500mM ammonium formate, pH 3. A starting elution gradient of 20% A, 30% B, 49% C and 1% D was linearly changed to 4% A, 14% B, 81% C and 1 % D for4.5 min, followed by isocratic elution at 4% A,14% B, 81% C and 1%D for 2.1 min and column equilibration with initial conditions for 1.4 min.

#### Mass Spectrometry Data Acquisition

Mass spectrometry data was acquired using ‘sensitivity’ mode in positive and negative electrospray ionization mode within 100-2000 Da. For the electrospray acquisition, the capillary voltage was set at 1.5kV (positive), 3.0kV (negative), sample cone voltage 30V, source temperature at 120°C, cone gas flow 50L/h and desolvation gas flow rate of 800L/h with scan time of 0.5 sec in continuum mode. Leucine Enkephalin; 556.2771 Da (positive) and 554.2615 Da (negative) was used for lockspray correction and scans were performed at 0.5sec. The injection volume for each sample was 3µL. The acquisition was carried out with instrument auto gain control to optimize instrument sensitivity over the samples acquisition time.

Data were processed using Progenesis QI (Nonlinear, Waters). Peak picking and retention time alignment of LC-MS and MSe data were performed using Progenesis QI software (Nonlinear, Waters). Data processing and peak annotations were performed using an in-house automated pipeline as previously described.^20–23^ Annotations were determined by matching accurate mass and retention times using customized libraries created from authentic standards and by matching experimental tandem mass spectrometry data against the NIST MSMS, LipidBlast or HMDB v3 theoretical fragmentations; for complex lipids retention time patterns characteristic of lipid subclasses was also considered. To correct for injection order drift, each feature was normalized using data from repeat injections of quality control samples collected every 10 injections throughout the run sequence. Measurement data were smoothed by Locally Weighted Scatterplot Smoothing (LOESS) signal correction (QC-RLSC) as previously described. Values are reported as ratios relative to the median of historical quality control reference samples run with every analytical batch for the given analyte^20–23^.

### Spatial Transcriptomic Analyses

Spatial transcriptomic data (Visium 10X platform) was derived from our recent foundational publication^17^. Single-cell data integration method to match and compare (scMC) pipeline was used for initial processing and clustering of the human IPMN ST dataset; to analyze spatially-resolved RNA-seq data, we used the Seurat package (version 3.2) implemented in R statistical software (version 4.3) (https://www.r-project.org/). ST data for the two *Kras;Gnas*^18^ samples was SCT normalized and merged. Spots in the Visium dataset were annotated as epilesional (Epi), meaning the areas covering the epithelial lining, juxtalesional (Juxta), corresponding to the adjacent microenvironment to the lesion, and perilesional (Peri), defined as an additional layer of stroma outside of the epithelium, as described.^19^

### Cell viability assay

Cell lines derived from murine IPMN-associated PDAC in the *Kras;Gnas* mice^18^ were seeded in a 96-well plate at a density of 2 × 10^4^ cells/well. Cell viability was determined using the CellTiter 96 Aqueous One Solution Cell Proliferation Assay (MTS) kit (Cat. #G3580, Promega). All experiments were performed in biological triplicates.

### Immunofluorescence

Murine IPMN-derived *Kras;Gnas* cells were fixed with 4% paraformaldehyde, permeabilized with 0.1% Triton X-100, and stained with CoraLite488 conjugated Tom20 antibodies (Cat. # CL48811802, Proteintech). Images were acquired in z-series on a spinning-disk confocal system.

### Immunoblotting

Whole cell lysate proteins were extracted using RIPA buffer (Cat. #89901, Pierce) containing complete protease inhibitor cocktails (Cat. #04693116001, Roche) and phosSTOP (Cat. #04906837001, Roche). The following antibodies were used for immunoblotting: p62 (Cat. # 39749, Cell Signaling Technology), Optineurin (Cat. #sc-166576, Santa Cruz Biotechnology), BNIP3L/Nix (Cat. # 12396, Cell Signaling Technology), FUNDC1 (Cat. # 49240, Cell Signaling Technology), LC3B (Cat. # 83506, Cell Signaling Technology), and β-actin (Cat. # 3700, Cell Signaling Technology).

### Animal study

Animal experiment protocols were approved by The University of Texas MD Anderson Cancer Center IRB and in accordance with the Guidelines for the Care and Use of Laboratory Animals published by the NIH (Bethesda, MD). Athymic nude (Cat. #002019, The Jackson Laboratory) were housed in specific pathogen free facilities. For subcutaneous allograft models of IPMN-associated PDAC, a total of 1 × 10^5^ 4861 cells were suspended in 100 µl 50% Matrigel Matrix (Corning)/Opti-MEM media and subcutaneously injected into 8-to 10-week-old mice. The mice were fed with water containing 0.2 mg/ml Doxycycline. For UGT8 inhibitor treatment, mice received UGT8-IN-1 (3mg/kg) via oral gavage every other day for 14 days starting from Day 7 post cancer cell inoculation. The tumor growth was monitored every 3 days.

### Gene expression datasets

Transcriptomic data and associated clinical information for The Cancer Genome Atlas (TCGA) PDAC dataset and the Clinical Proteomic Tumor Analysis Consortium (CPTAC) PDAC dataset^24^ was downloaded from cBioportal (https://www.cbioportal.org/).^25^ Single cell RNA-sequencing data for 6 patients with PDAC was retrieved from the Gene Expression Omnibus dataset (GSE212966)^26^. The R package “Seurat” was utilized for data preprocessing; principal component analysis (PCA)-based dimension reduction was performed and the FindCluster function applied to cluster cell types. The t-distributed stochastic neighbor embedding (t-SNE) method was used to visualize the distinct cell clusters. The Findmarkers function was used to identify marker genes in each clusters and the cell clusters were annotated based on highly expressed genes and reported canonical cellular markers^27^.

### Statistical Analyses

Statistical significance was determined using Dunn’s multiple comparison tests and Bonferroni-adjusted p-values are reported unless otherwise specified. For in vivo experiments, statistical significance was determined using repeated measures two-way ANOVA and 2-sided p-values reported for the treatment effect.

Unsupervised Hierarchical Clustering heatmaps of MALDI-MS-derived ion features for IPMN tissues were generated using R statistical software (V4.3.0) (https://www.r-project.org/). Univariable and multivariable Cox Proportional hazard models were used to evaluate associations between tumoral UGT8 mRNA expression stratified into high (> median) and low (<= median) and overall survival. To test for the proportionality of hazard assumption of a Cox regression, we used the method of Patricia and Grambsch.^28^ Kaplan-Meier curves were generated in R statistical software. Log-rank (Mantel-Cox) tests were used to compare survival curves and 2-sided p-values reported.

## RESULTS

### MALDI-MS imaging reveals selective localization of sulfatide species to the IPMN and PDAC epithelium

We applied matrix-assisted laser desorption/ionization (MALDI)-mass spectrometry (MS) to perform an unbiased spatial characterization of lipid profiles on frozen tissue sections from resected human IPMN cases. The specimen set consisted of 23 IPMN tissue sections with either low-grade dysplasia (LG) (n=15), high-grade dysplasia (HG) (n=4), or with an associated PDAC (referred to hereon as IPMN/PDAC; n=4) (**Table S1**). Various lipid species, including lysophospholipids, glycerophospholipids, and sphingolipids, were putatively identified in the MALDI-MS imaging mass spectra from the IPMN specimens based on exact mass values matched to the LipidMaps database (**Fig. S1 and Table S2**). MALDI mass spectra obtained from stromal areas, adjacent normal pancreas tissue, including ductal and acinar cells, as well as regions of the neoplastic epithelium were extracted, resulting in 1,168 lipid features that were used for further statistical analyses. Unsupervised hierarchical clustering heatmaps revealed the lipidome of IPMN to be distinct from normal epithelium or exocrine pancreas (**Figure 1A**), with 898 of the 1,168 lipid features achieving statistical significance (FDR-adjusted Wilcoxon rank sum test 2-sided p<0.05) (**Table S2**).

**Figure 1.**
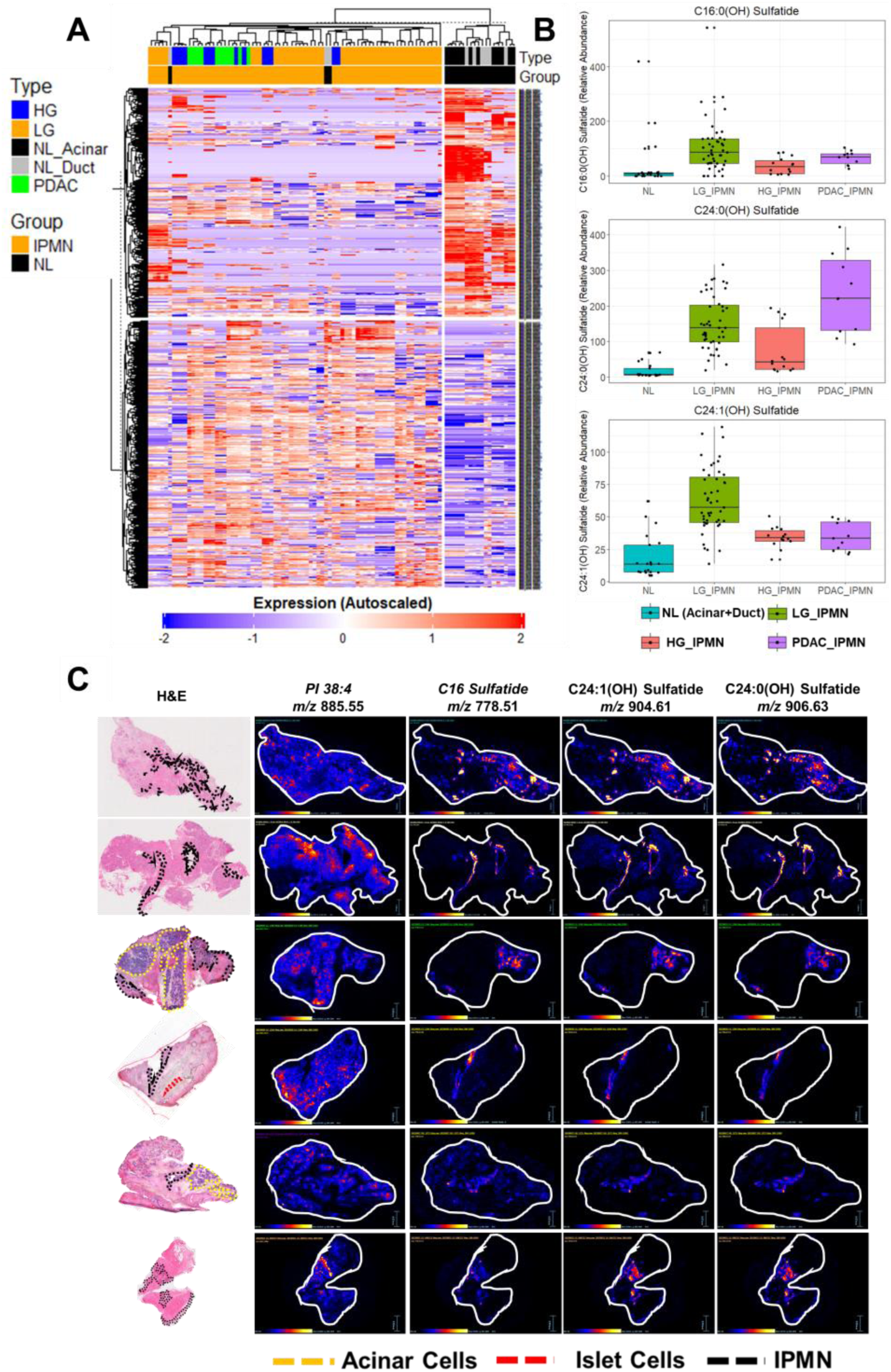
MALDI-MS imaging of PDAC and IPMN. **A)** Unsupervised hierarchical clustering heatmaps of lipid features extract from the neoplastic epithelium and from stromal areas and adjacent normal pancreas tissue. **B)** Box and whisker plots for C16:0, C24:0(OH), and C24:1(OH) sulfatides normal pancreas tissue (NL-including ductal and acinar cells), as well as regions of the neoplastic epithelium. **C)** Representative MALDI-MS images showing localization of selected sulfatide species in a PDAC (top) and LG IPMN (bottom) tissue sample. Optical image for a serial tissue section stained with hematoxylin and eosin (H&E) is provided with the PDAC and IPMN areas outlined. A MALDI-MS image for the ion at *m/z* 885.55, which corresponds to a glycerophosphoinositol 38:4 is shown for both samples as comparison. A white outline showing the edge of the tissue is drawn on top of the MALDI-MS images.

Among the top differential lipids elevated in IPMN were several sulfatide species including C16 sulfatide (*m/z* 778.51), C24:0(OH) sulfatide (*m/z* 906.63), and C24:1(OH) sulfatide (*m/z* 904.61) (**Fig. 1B; Table S2**). Remarkably, evaluation of MALDI-MS images and corresponding tissue histology showed a prominent and selective enrichment of the C16, C24:0(OH), and C24:1(OH) sulfatides in areas outlining the IPMN and cancer epithelium (**Fig. 1C**). Confirmation of sulfatide annotations were performed by comparing the exact mass and isotope patterns between the tissue MALDI mass spectrum to that obtained using dried sulfatide standards (**Fig. S2**).

### Spatial transcriptomic analyses of IPMN tissues supports increased sulfatide metabolism in IPMN

Using a recently published dataset of spatial transcriptomic (ST) data derived from the Visium 10X platform for serial sections derived from the same tissues evaluated via MALDI-MS imaging, we investigated co-occurrence of RNA transcripts corresponding to enzymes involved in sulfatide metabolism.^17^ Spots in the Visium dataset were previously annotated as epilesional (Epi), defined as the areas covering the epithelial lining, juxtalesional (Juxta), corresponding to the adjacent microenvironment to the lesion, and perilesional (Peri), defined as an additional layer of stroma outside of the epithelium. ST analyses revealed RNA transcript levels of *ceramide galactosyltransferase* (*CGT*, also known as *UGT8*) and *galactose-3-O-sulfotransferase* (*CST*, also known as *Gal3St1*), enzymes that, respectively, catalyze the addition of galactose from UDP-galactose to ceramide^29^ and the *O*-sulfation of the galactose residue on galactosylceramide^30^, to be highly enriched in epilesional spots from the Visium dataset of IPMN and PDAC samples (**Fig. 2A-C; Fig. S2**). In comparision, transcript levels of *UDP-glucose ceramide glucosyltransferase* (*UGCG*) as well as *lactosylceramide synthase* (encoded by *beta-1,4-galactosyltransferase 5* (B4GALT5)), which encode for enzymes involved in synthesis of glucosylceramides and lactosylceramides, respectively, (**Fig. S3**), tended to be more broadly expressed in other areas of the tissues such as the stroma and lymphocyte aggregates (**Fig. S4**).

**Figure 2.**
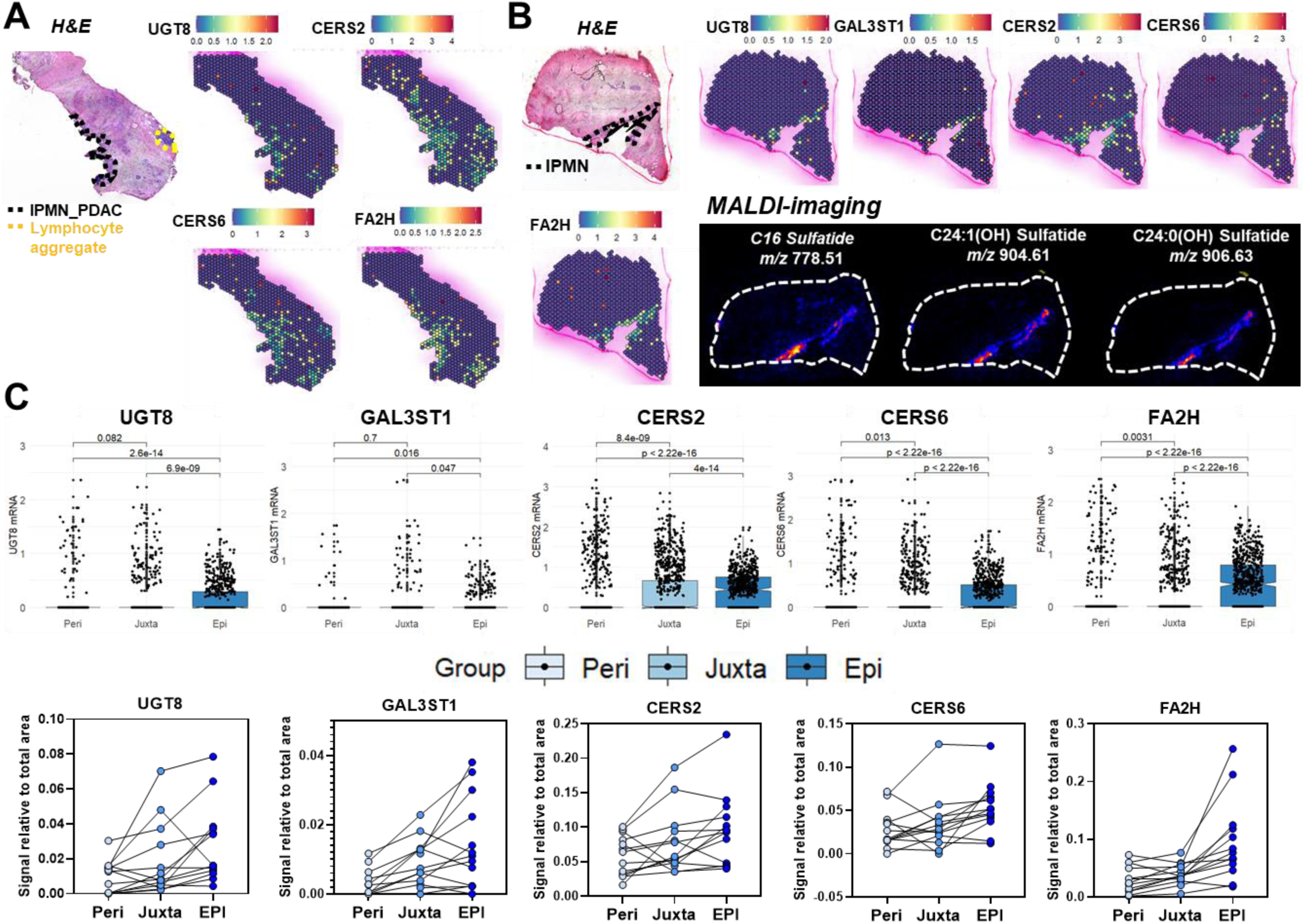
Spatially resolved transcriptomics of IPMN and PDAC tissue samples. **A)** Spatial transcriptomics demonstrates enrichment of UGT8, CERS1, CERS6, and FA2H transcript levels in IPMN with associated adenocarcinoma. **B)** Spatial transcriptomics and MALDI-MS images obtained from the same sample using serial sections demonstrated enrichment of UGT8, GAL3ST1, CERS2, CERS6, and FA2H transcript levels to be co-localized in the IPMN epithelium with sulfatides. The IPMN area is outlined in black. **C)** Boxplot showing gene expression levels for UGT8, GAL3ST1, CERS2, and FA2H for spots in epilesional (Epi), juxtalesional (Juxta), and perilesional (Peri) areas. Statistical significance was determined by Dunn’s multiple comparison test and adjusted P-values are reported. Statistical significance was determined by Dunn’s multiple comparison test and adjusted P-values are reported. Dot plots beneath represent average transcript levels normalized to total area for Epi, Juxta, and Peri regions. Connecting lines represent corresponding regions from the same tissue sample.

Concomitant to the increases in *UGT8* and *GAL3ST1* were elevated transcript levels of fatty acid 2-hydroxylase (*FA2H*), ceramide synthase 2 (*CERS2*) and *CERS6*, which encode for enzymes involved in free fatty acid hydroxylation and *de novo* synthesis of long-chain (C22-C24) and palmitoyl-chain (C16) ceramides, respectively (**Fig. 2A-C; Fig. S5A**). In contrast, *CERS4* (C18-C20) and *CERS5 (*C14-C16) tended to be lower in IPMN epithelium; *CERS1* and *CERS3* were not differentially expressed (**Fig. S5A**). These data collectively support higher activity of the ceramide synthases with preferences towards longer acyl chain lengths. To this end, direct comparison of spatial MALDI-MS and ST data from the same IPMN lesion demonstrated that mRNA levels of *FA2H*, *CERS2, UGT8*, and *GAL3ST1* in IPMN epithelium strongly coincided with enrichment of C16, C24:0(OH), and C24:1(OH) sulfatides (**Fig. 2B**). *UGT8*, *GAL3ST1*, *CERS2*, *CERS6*, and *FA2H* were not significantly differential between LG and high-risk (HG + IPMNPDAC) IPMN, suggesting that upregulation of these enzymes occurs during early states of neoplastic development and persists during progression (**Fig. S5B**).

### Cross-species MALDI-MS imaging reveals long-chain hydroxylated sulfatide to be highly specific to the neoplastic epithelium in IPMNs

To further evaluate the relationship between enhanced sulfatide metabolism and progression of IPMN towards malignancy, we performed MALDI-MS imaging of tissue sections from resected cystic lesions from p48-Cre; LSL-KrasG12D; Rosa26R-LSL-rtTA-TetO-GnasR201C mice (hereon referred to as *Kras;Gnas* mice)^31^. Consistent with our findings in human IPMN, spatial distribution of the C24:1(OH) and C24:0(OH) sulfatide outlined the epithelial cystic lining, as shown in **Fig. 3** for a neoplasm collected from a *Kras;Gnas* mice after 25 weeks on doxycycline diet. Of relevance, C24:1(OH) and C24:0(OH) sulfatide levels were more prominent in the cystic areas presenting with a higher grade of dysplasia (**Fig. 3A-Panel A**) compared to neighboring smaller cysts with lower grade of dysplasia (**Fig. 3A-Panel B**). Visium spatial transcriptomics analysis of pancreatic tissue samples collected from the *Kras;Gnas* mice after 25 weeks on doxycycline diet also yielded detectable transcripts for sulfatide-related enzyme transcripts including *Ugt8a*, *Gal3st1*, and *Fa2h*. As shown in *Fig. 3C*, elevated levels of *Gal3st1* and *Fa2h*, and to a lesser extent *Ugt8*, were observed in the cystic lesion areas (**Fig. 3B**).

**Figure 3.**
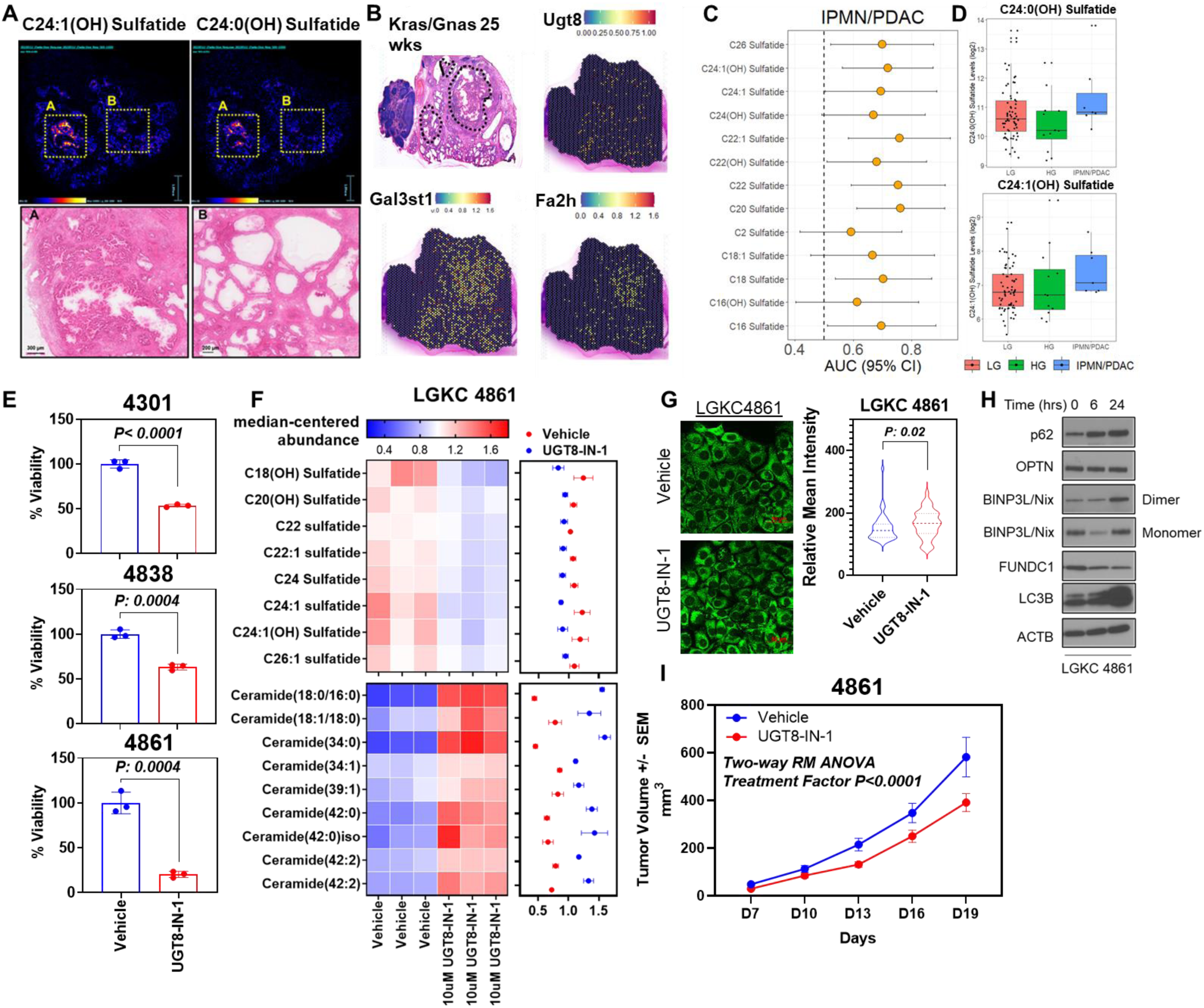
Enhanced sulfatide metabolism is a metabolic feature that is associated with IPMN development in *Kras;Gnas* mice. **A)** (Top) MALDI-MS images showing distribution of C24:1(OH) and C24:0(OH) ST in a *Kras;Gnas* mouse tissue section harvested after 25 weeks on doxycycline diet. (Bottom) H&E staining with higher magnification for two regions: (A) high dysplasia, (B) low dysplasia. **B)** Spatial transcriptomics from the same sample using serial sections as in panel (A) demonstrated enrichment of UGT8, GAL3ST1, and FA2H transcript levels in IPMN cysts from *Kras;Gnas* mice. **C)** Dot plot demonstrating individual classifier performance (AUC (95% confidence intervals)) for individual sulfatides detected in cystic fluid by LCMS for distinguishing IPMN/PDAC from LG IPMN. **D)** Box and whisker plots showing levels of C24:0(OH) and C24:1(OH) sulfates in cystic fluid of patients with LG IPMN (n=68), HG IPMN (n=12), or IPMN/PDAC (n= 7). **E)** In vitro toxicity assays (MTS Assay) for three independent mouse IPMN cell lines (4301, 4838, and 4861) treated with vehicle control or the UGT8 inhibitors UGT8-IN-1 (10 µM). Experiments were performed in technical triplicates. Statistical significance was determined by 2-sided Student T-test. F) Heatmap depicting median-centered abundances of significantly (2-sided Student T-test p< 0.05) differential sulfatides (top panel) and ceramides (bottom panel) in murine 4861 IPMN cells following 6-hour treatment with 10μM UGT8-IN-1 compared to vehicle control. Dot plots to the right indicate median center abundances +/− standard deviation of respective sulfatide and ceramides species. Each experiment consisted of 3 biological replicates. **G)** Representative images of Tom20^+^ GFP 4861 cells following 6-hour treatment with 10μM UGT8-IN-1 or vehicle (DMSO) control. All images were captured via confocal and widefield microscopy using live cell imaging techniques. Scale bars represent 50 µm; digital image acquisition parameters and look up table mappings were uniformly set for all images within each respective. Quantitative values (relative mean intensities) are provided to the right of the images. **H)** Immunoblots for mitophagy-related proteins in murine 4861 cells following 6-hour treatment with 10μM UGT8-IN-1 or vehicle (DMSO) control. **I)** Tumor volume in LGKC4861 (p48-Cre; LSL-KrasG12D; Rosa26R-LSL-rtTA-TetO-GnasR201C)-tumor bearing mice following treatment with saline control or 3mg/kg UGT8-IN-1. Tumor volume was measured every 3 days. Statistical significance was determined by 2-way repeated measures ANOVA and treatment factor 2-sided p-value reported.

### Long-chain hydroxylated sulfatides are elevated in cystic fluid of patients with malignant IPMN

We additionally performed lipidomic analyses of cystic fluid collected from an independent cohort of 89 patients with IPMN. A total of 14 sulfatide species were detected and quantified in cystic fluid (**Table S3**). Of the 14 quantified sulfatide species, 12 were elevated in cystic fluid of patients harboring IPMN/PDAC, with individual predictive performance estimates (AUC) ranging from 0.61-0.76 (**Fig. 3C; Fig. S6; Table S3**). Importantly, both C24:0(OH) and C24:1(OH) sulfatides were elevated in patients with malignant IPMN compared to patients with LG IPMN (**Fig. 3D**). Taken together with our findings of elevated sulfatides in cystic fluid coupled with MALDI-MS analyses of IPMN tissues supports elevated sulfatide biogenesis as an early metabolic manifestation in cystic neoplasms that persists into malignancy.

### Sulfatide metabolism represents a therapeutically targetable metabolic vulnerability in IPMN

We further assessed the therapeutic potential of targeting sulfatide metabolism in the context of IPMN. Using mass spectrometry, we first confirmed that sulfatides, including the 24:0(OH) and 24:1(OH) sulfatide, are readily detectable in multiple IPMN-derived murine PDAC cell lines that were generated from pancreata of *Kras;Gnas* mice^18^ (**Fig. S7**). Treatment of three independent doxycycline (100ng/mL)-induced *Kras;Gnas* cell lines (4301, 4838, and 4861) with 10μM UGT8-IN-1 resulted in marked reductions (Student T-test 2-sided p< 0.05) in cell viability compared to vehicle control (**Fig. 3E**). Lipidomic analyses of *Kras;Gnas* 4861 total cell extracts following short-term treatment (6 hours) with UGT8-IN-1 revealed profound changes in lipid composition, including significant (2-sided T-test p<0.05) reductions in several sulfatide species, such as 24:1(OH) (**Fig. 3F; Table S4**). These changes were accompanied by a concomitant increase in several ceramides, suggesting a block in downstream sulfatide biosynthesis (**Fig. 3F; Table S4**).

Ceramides are reported to exert anti-cancer effects through induction of lethal mitophagy^23,32^. Assessment of mitochondrial morphology in *Kras;Gnas* 4861 cells following 6-hour treatment with UGT8-IN-1 indicated accumulation of Tom20-GFP^+^ mitochondria (Wilcoxon rank sum test 2-sided P: 0.02). Immunoblots demonstrated time-dependent increases in PINK1-mediated mitophagy markers, p62 and optineurin (OPN), as well as the autophagy/mitophagy marker LC3B following UGT8-IN-1 treatment, consistent with induction of mitophagy (**Fig. 3H**).

To assess for in vivo efficacy of UGT8-IN-1, *Kras;Gnas* 4861 cells (1*10^5^ cells per mouse) were implanted in 8-week-old athymic nude mice (N= 6 per group; 3 males and 3 females) subcutaneously, and the mice were administered doxycycline (200 ug/ml) supplemented in drinking water. The UGT8 inhibitor UGT8-IN-1 or vehicle control (saline) was orally administrated every other day (3 mg/kg) for 2-weeks. Treatment of 4861-tumor bearing mice with UGT8-IN-1 led to attenuation of tumor growth (Two-way Repeated Measures ANOVA treatment effect 2-sided p<0.0001) (**Fig. 3I**). Taken together, these results highlight the therapeutic potential of targeting sulfatide metabolism in IPMNs and PDAC lesions.

### Long-chain hydroxylated sulfatides are prevalent in PDAC and elevated *UGT8* mRNA expression is prognostic for poor overall survival

To assess for broader relevance of our findings in the context of PDAC, we performed MALDI-MS imaging of an additional cohort of PDAC tissues obtained from nine cases without an associated IPMN (**Table S1**). Representative MALDI-MS imaging of one PDAC sample containing areas of perineural invasion is illustrated (**Fig. 4A**), demonstrating selective enrichment of sulfatides in the cancerous tissue, colocalized with the shape of the cancer region surrounding the nerve. **Fig. 4B** further demonstrates enrichment of C24:0(OH) and C24:1(OH) sulfatides in PDAC compared to normal tissues or tissues from patients with chronic pancreatitis.

**Figure 4.**
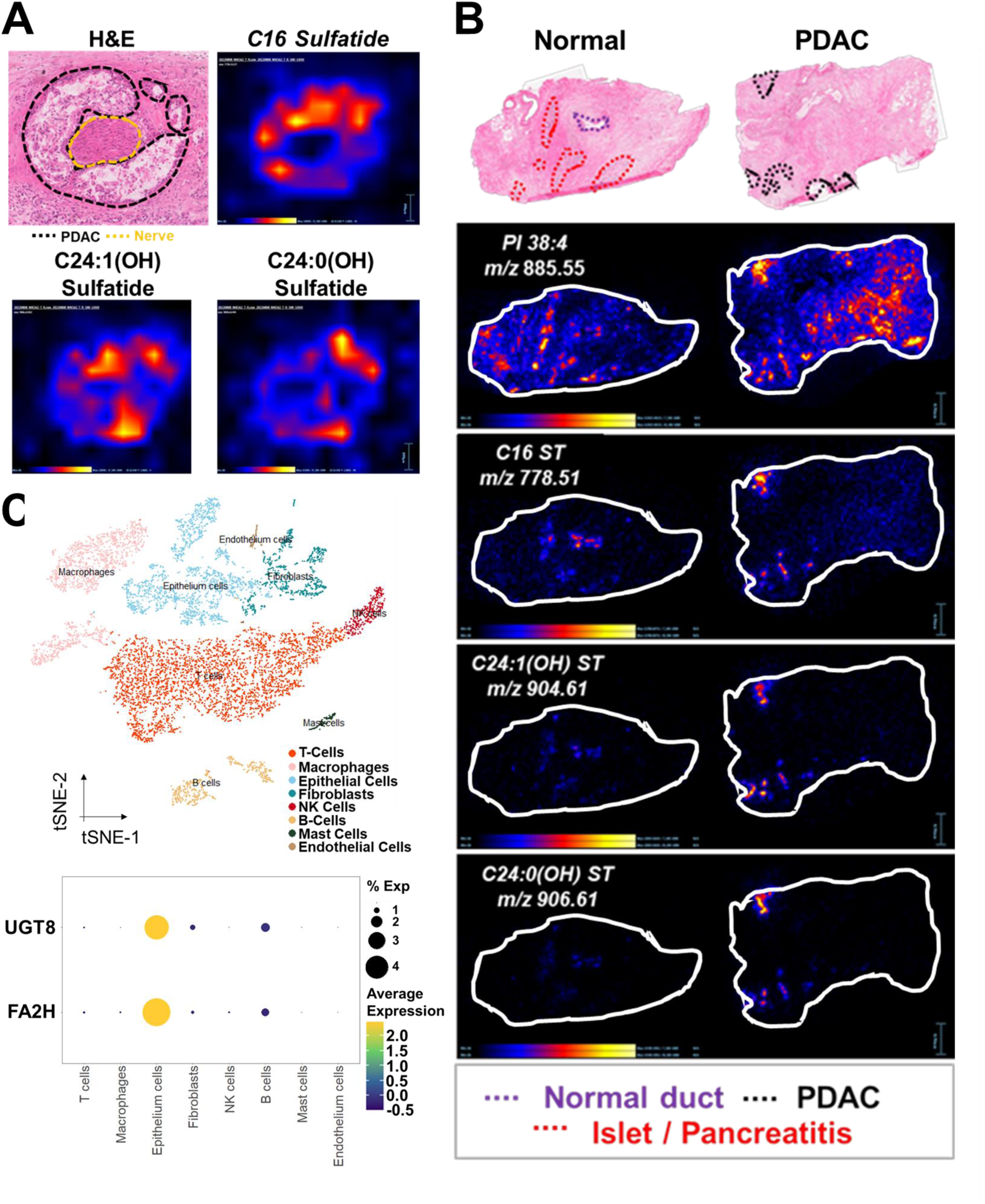
Long-chain hydroxylated sulfatides are selectively enriched in PDAC. **A)** MALDI-MS imaging showing localization of the C16, C24:0(OH), and C24:1(OH) sulfatides to PDAC lesion surrounding a nerve ending. **B)** MALDI-MS images showing distribution of C16, C24:0(OH), and C24:1(OH) sulfatides in PDAC tissues and tissues from patients with chronic pancreatitis. Glycerophosphoinositol 38:4 is shown for both samples as comparison. **C)** Single cell RNA-sequencing data for 6 patients with PDAC was retrieved from the Gene Expression Omnibus dataset GSE212966. tSNE visualization of 8 cell-type clusters are provided (top panel). Dot plots demonstrate high percentage and expression of UGT8 and FA2H transcripts in epithelial cells (bottom panel).

Since matched ST data was not available on these nine independent cases, we compared expression levels of transcripts encoding for sulfatide-biosynthesis enzymes in human PDAC tumors from The Cancer Genome Atlas (TCGA), compared with normal PDAC tissues from The Genotype-Tissue Expression (GTEX) project. These analyses also revealed transcript levels of *UGT8*, *GAL3ST1*, *CERS2*, *CERS6*, and *FA2H* to be highly (Wilcoxon Rank Sum test 2-sided p<0.0001) elevated in PDAC tissues (**Fig. S8**). Single cell RNAseq data for 6 PDAC tumors from the Gene Expression Omnibus dataset (GSE212966)^26^ further demonstrated transcript levels of *UGT8* and *FA2H* to be particularly enriched in malignant epithelium cells compared to other cell types (**Fig. 4C**).

We assessed for association between *UGT8* mRNA expression in PDAC tumors with overall survival in the Clinical Proteomic Tumor Analysis Consortium (CPTAC) dataset^33^. High tumoral *UGT8* mRNA expression and protein expression of UGT8 (> median) was associated with worse overall survival in PDAC (**Fig. S9**). Multivariable Cox proportional hazard models, considering age, sex and tumor stage as co-variables, similarly showed that elevated tumoral *UGT8* mRNA (Hazard Ratio (95% CI): 1.80 (1.13-2.89)) levels were a statistically significant independent prognostic indicators of poor overall survival in PDAC (**Table 1**). Assumption of the proportionality of hazard assumption of the Cox regression model were met in our analyses. Similar findings were found in The Genome Cancer Atlas (TCGA) PDAC transcriptomic dataset (Multivariate Cox Proportional Hazard Ratio (95% CI): 1.48 (0.97-2.87)) (**Fig. S9; Table 1**).

**Table 1.**
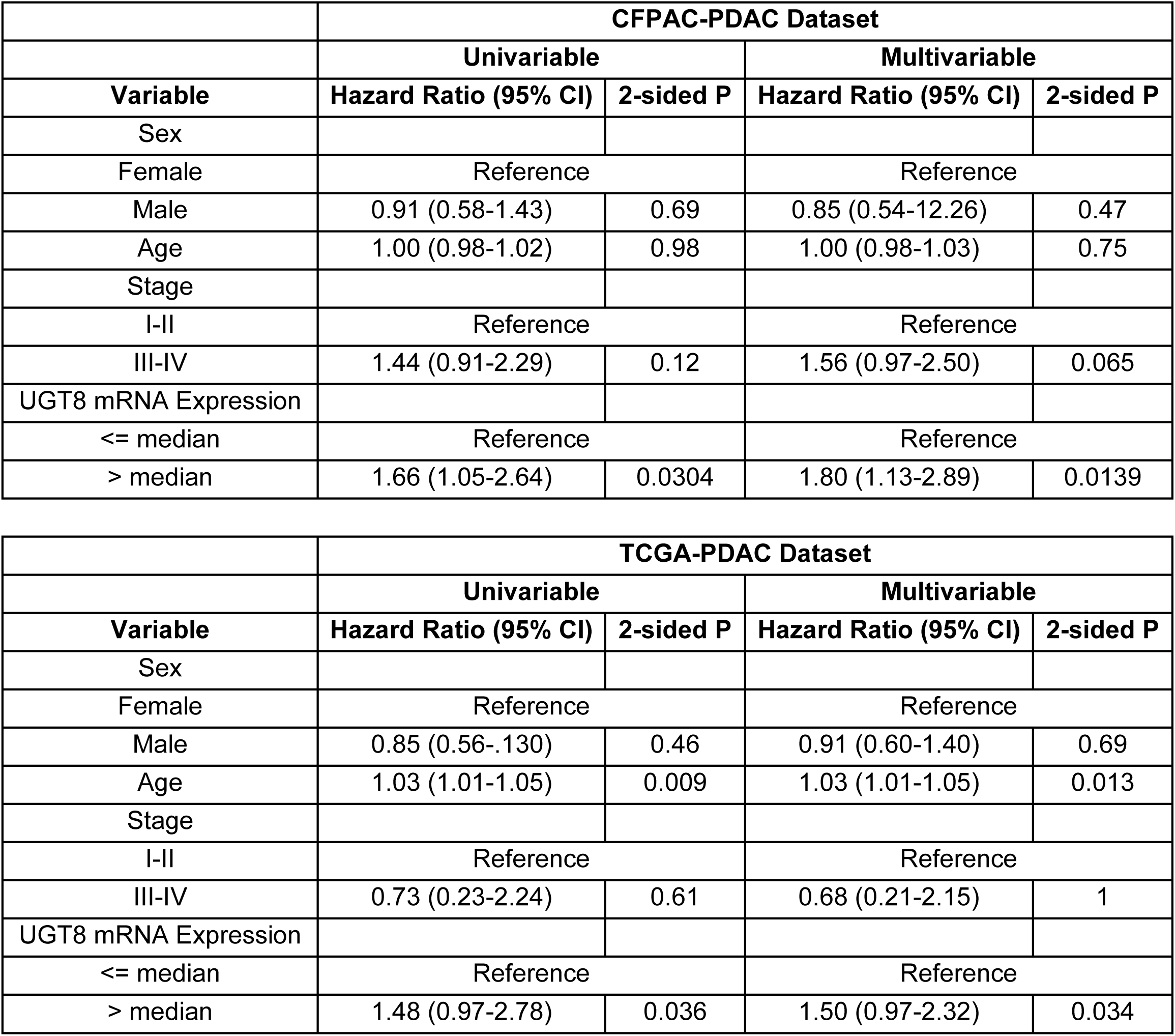
Cox proportional hazard models illustrating association between tumoral UGT8 mRNA levels and overall survival in PDAC patients in the CFPAC and TCGA PDAC datasets.

## Discussion

Our integrated cross-species spatial-omics analyses have led to the discovery of specific enrichment of hydroxylated long chain sulfatides in human and murine epithelium in the context of IPMN and that enhanced sulfatide metabolism is associated with IPMN and persists to malignancy. Importantly, we provide evidence that sulfatide metabolism represents an actionable a metabolic vulnerability with therapeutic potential.

To-date, the cell autonomous role(s) of sulfatide in the context of cancer development and progression are poorly understood. Sulfatides represent a subtype of sphingolipid typically found in myelin sheath. Sulfatides are variable in structure, comprising different lengths of acyl chain and ceramide moiety, which can be hydroxylated, as well as other sphingolipids^30^. Sulfatide synthesis begins in the endoplasmic reticulum by the addition of galactose from UDP-galactose to ceramide, which is catalyzed by the ceramide galactosyltransferase (CGT, also known as UGT8)^29^ followed by *O*-sulfation of the galactose residue on galactosylceramide via galactose-3-O-sulfotransferase (CST, also known as Gal3St1) that occurs in the Golgi apparatus^30^. The 3-*O*-sulfate group of sulfatide can be hydrolyzed by arylsulfatase A (ASA; also referred to as ARSA)^30^.

A high level of sulfatide is a frequently observed phenomenon in cancer cell lines and tumor tissues, including lung adenocarcinoma^34^, gastric, kidney, ovarian, breast, and colorectal cancer and elevated sulfatide level in ovarian and colorectal tumors is associated with poor survival^35,36^. Yet, mechanistic understanding of deregulated sulfatide in the context of early neoplasia has not been explored. Prior studies suggest that sulfatide is a native ligand for L- and P-selectin, which plays a role in facilitating disease progression, metastasis^37^, and immune modulation. For example, binding of sulfatide to L-selectin was shown to up-regulate expression of the chemokine co-receptor CXCR4 surface expression CD4+ T cells^38^. Other chemokines including CCL2, CCL3, CCL4, CXCL8, and CXCL12 have also been reported to selectively bind sulfatide^39^, which may promote migration of lymphocytes to cancer cells abundance in sulfatide^38^. Moreover, sulfatide has been shown to directly bind receptors on macrophages, resulting in enhanced TGF-beta1 and IL-6 secretion, and P-selectin expression^40^. Treatment of antigen-presenting cells with C24:1 sulfatide increased expression of indoleamine 2,3-dioxygenase 1 (IDO1)^41^, an rate-limiting enzyme in the kynurenine pathway that is involved in tumor immune suppression^42,43^. Collectively, these findings suggest a role of sulfatide in modulating the immune response, an aspect that is currently under investigation, although outside of the scope of the current study.

ST analyses identified *UGT8* and *FA2H* to be particularly elevated in neoplastic epithelium and to co-occur with enrichment of sulfatides. Single-cell RNA sequencing data from PDAC tumors similarly revealed *UGT8* and *FA2H* to be highly elevated in epithelial cells compared to other cell types in the tumor microenvironment. Elevated tumoral UGT8 expression in basal-like breast cancer is associated with poor overall survival. Mechanistically, Sox10-mediated upregulation of UGT8 induced the sulfatide biosynthetic pathway, leading to induction of integrin aVb5-mediated signaling to enhance cell migration and invasion. Knockdown of UGT8 attenuated migration and invasion of TNBC cell lines, which could be restored through supplementation with sulfatides^33^. High *UGT8* and *FA2H* transcript levels also correlates with elevated levels of hydroxylated hexosylceramides in lung adenocarcinoma^44^. Enrichment of FA2H in subpopulation of esophageal squamous cell carcinoma (ESCC) cells is associated with high metastatic potential and poor overall survival of ESCC patients. FA2H expression was found to be transcriptionally induced through a TNFa-FOXC2-FA2H signaling axis^45^. These studies indicate additional cancer cell intrinsic properties of enhanced sulfatide metabolism yielding pro-tumoral effects.

In conclusion, our study identifies a novel metabolic adaptation of enhanced sulfatide biosynthesis as an early metabolic manifestation of precursors in the pancreas and that presents as a potential interception strategy for attenuating or preventing malignant progression.

## ACKNOWLEDGEMENTS

A.M. is supported by the Sheikh Khalifa bin Zayed Foundation, the MD Anderson Pancreatic Cancer Moonshot, The University of Texas MD Anderson Cancer Center Moon Shots Program™, and NCI P50CA221707, U01CA200468, U54CA274371. M.S. is supported by TRIUMPH Fellowship in the CPRIT Training Program (RP210028). M.Y.S., J.Z., C.M.S., S.H., and J.F.F. are supported by NCI grant U01CA239522.

## CONFLICTS OF INTEREST

AM is listed as an inventor on a patent that has been licensed by Johns Hopkins University to Thrive Earlier Detection and serves as a consultant for Tezcat Biosciences. JF, SH, MS and AM have filed a patent related to the findings in this manuscript that is pending.

---

**Table S1.** Patient and tumor characteristics for tissues analyzed by MALDI-MS.

| Sample | ST | MALDI-MS | Normal Adjacent Tissue MALDI-MS | Gender | Age | Diagnosis | Grade, Pathology | Location | TNM | Stage | Neoadjuvant Treatment |
| --- | --- | --- | --- | --- | --- | --- | --- | --- | --- | --- | --- |
| LG_1 | y | y | n | F | 75 | IPMN | LG IPMN | head | - | - | n |
| LG_2 | y | y | n | M | 76 | IPMN | LG IPMN | head | - | - | n |
| LG_3 | y | y | n | M | 56 | IPMN | LG IPMN | head, body | - | - | n |
| LG_4 | y | y | n | F | 67 | IPMN | LG IPMN | head | - | - | n |
| LG_5 | y | y | n | F | 76 | IPMN | LG IPMN | body/tail | - | - | n |
| LG_6 | y | y | n | F | 51 | IPMN | LG IPMN | body/tail | - | - | n |
| LG_7 | y | y | n | M | 76 | IPMN | LG IPMN | body/tail | - | - | n |
| LG_8 | n | y | n | F | 72 | IPMN | LG IPMN | body/tail | - | - | n |
| LG_9 | n | y | n | M | 72 | IPMN | LG IPMN | head | - | - | n |
| LG_10 | n | y | n | F | 76 | IPMN | LG IPMN | body/tail | - | - | n |
| LG_11 | n | y | n | F | 72 | IPMN | LG IPMN | head, body, tail | - | - | n |
| LG_12 | n | y | n | F | 68 | IPMN | LG lesion<br>from HG<br>IPMN<br>case | body/tail | - | - | n |
| LG_13 | n | y | n | F | 72 | IPMN | LG IPMN | head,<br>body | - | - | n |
| LG_14 | n | y | y | F | 73 | IPMN | LG lesion<br>from HG<br>IPMN | body/tail | - | - | n |
| LG_15 | n | y | n | M | 74 | IPMN | LG lesion<br>from HG<br>IPMN | head | - | - | n |
| HG_1 | y | y | n | M | 72 | IPMN | HG IPMN | head | - | - | n |
| HG_2 | y | y | n | F | 89 | IPMN | HG IPMN | head | - | - | n |
| HG_3 | y | y | n | M | 62 | IPMN | HG IPMN | head | - | - | n |
| HG_4 | n | y | n | F | 67 | IPMN | HG IPMN | head | - | - | n |
| PDAC_1 | y | y | n | F | 76 | IPMN/PDAC | PDAC with<br>co- occurring<br>IPMN | head | - | - | n |
| PDAC_2 | y | y | n | M | 85 | IPMN/PDAC | PDAC with<br>co-<br>occurring<br>IPMN | head,<br>uncinate | - | - | n |
| PDAC_4 | n | y | n | M | 84 | IPMN/PDAC | PDAC<br>arising in<br>IPMN | body | pT2, pN0,<br>cM0, G2 | IB | n |
| PDAC_10 | n | y | n | F | 74 | IPMN/PDAC | PDAC with<br>HG IPMN | head,<br>uncinate | ypT3,<br>ypN0,<br>cM0, G2 | III | CT/RT |
| PDAC_5 | n | y | n | M | 69 | PDAC | PDAC | head | ypT2,<br>ypN2,<br>cM0, G3 | III | CT/RT |
| PDAC_6 | n | y | n | M | 66 | PDAC | PDAC | head | ypT2,<br>ypN2,<br>cM0, G3 | III | CT |
| PDAC_7 | n | y | y | M | 69 | PDAC | PDAC | head | ypT2,<br>ypN1,<br>cM0, G3 | IIB | CT |
| PDAC_8 | n | y | y | F | 62 | PDAC | PDAC | neck | ypT1c,<br>ypN0,<br>cM0, G2 | IA | CT/RT |
| PDAC_9 | n | y | n | F | 71 | PDAC | PDAC | head | ypT2,<br>ypN1,<br>cM0, G2 | IIB | CT |
| PDAC_11 | n | y | n | F | 75 | PDAC | PDAC | head | ypT2,<br>ypN2,<br>cM0, G2 | III | CT |
| PDAC_12 | n | y | y | M | 74 | PDAC | PDAC | body | ypT2,<br>ypN1,<br>cM0, G2 | IIB | CT |
| PDAC_13 | n | y | y | M | 67 | PDAC | PDAC | head | ypT3,<br>ypN1,<br>cM0, G3 | IIB | CT |
| PDAC_14 | n | y | n | M | 63 | PDAC | PDAC | body | ypT2,<br>ypN1,<br>cM0, G3 | IIB | CT |

**Figure S1.**
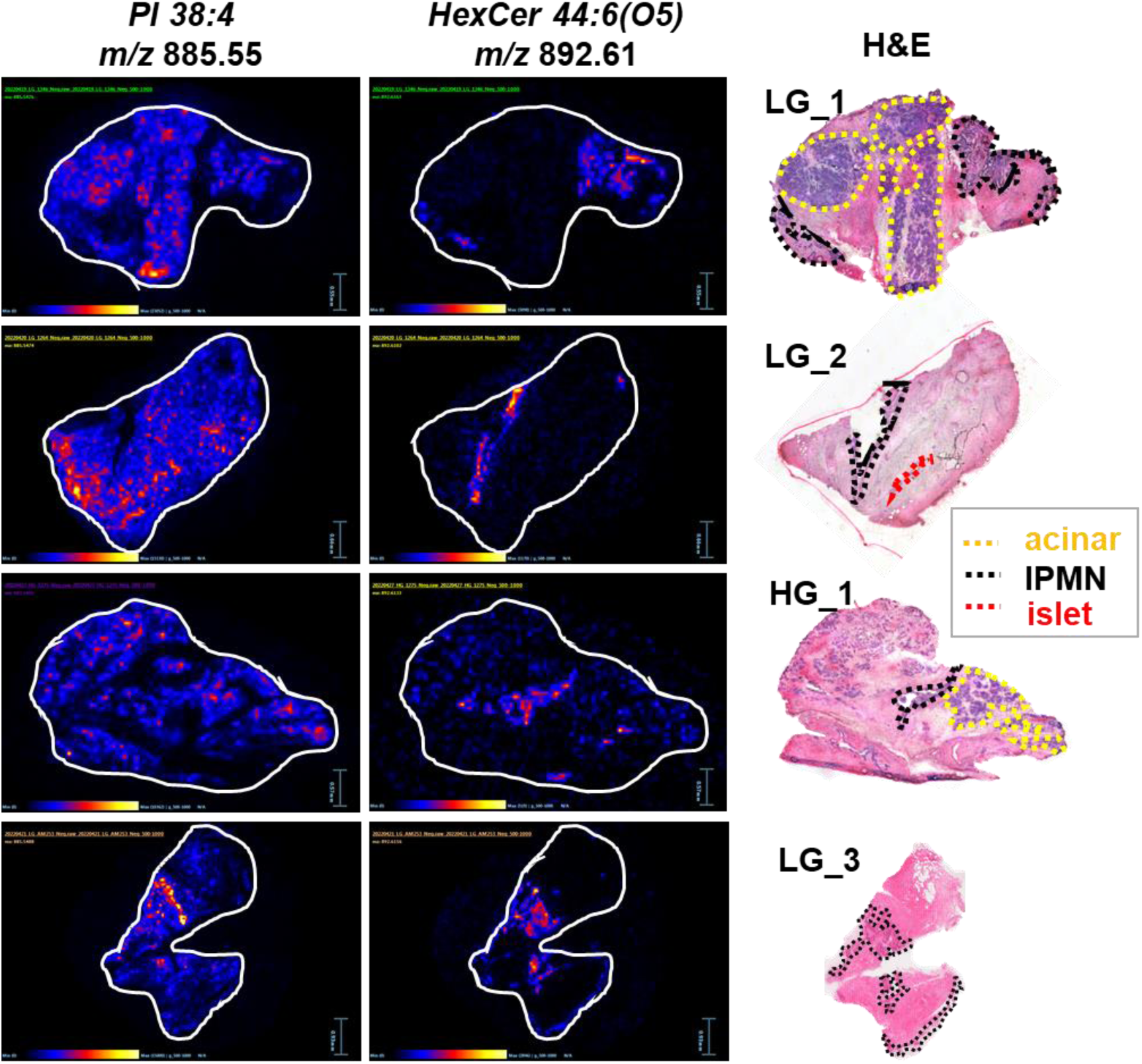
MALDI-MS Imaging reveals enrichment of cerebroside species in IPMN. Representative MALDI-MS images of hydroxylated hexosylceramide(44:6;O5) in tissue with high-grade (HG) and low-grade (LG) IPMN. A MALDI-MS image for glycerophosphoinositol (PI:38:4) is shown for comparison. The IPMN area is outlined in black, while a region containing lymphocyte aggregates or islets are outlined in yellow and red, respectively.

**Figure S2.**
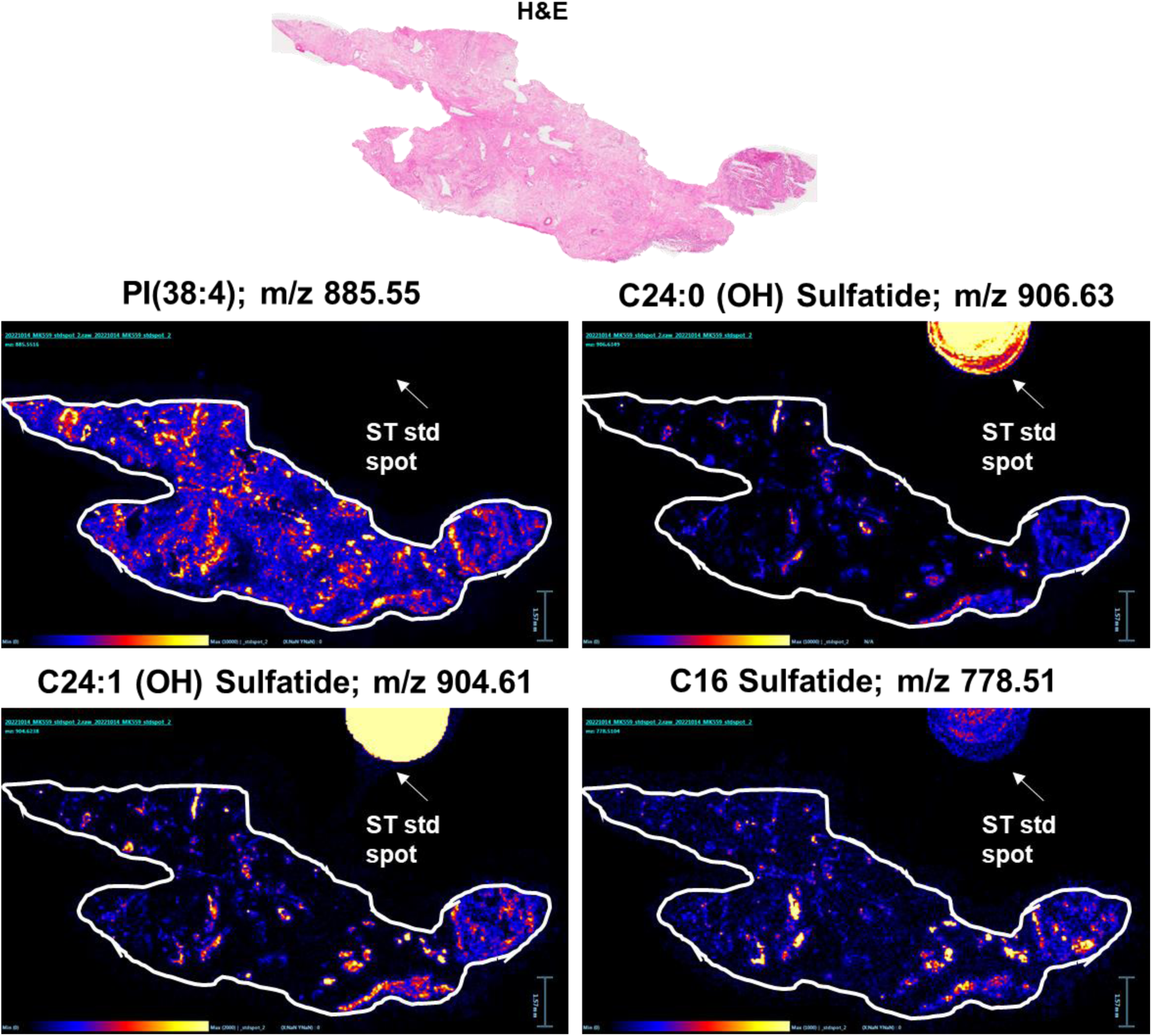
MALDI-MS imaging of dried sulfatide standards. Representative MALDI-MS images of. C16, C24:0(OH), and C24:1(OH) sulfatides in IPMN tissues as well as a dried sulfatide standard (ST Std) spot. A MALDI-MS image for glycerophosphoinositol (PI:38:4) is shown for comparison.

**Figure S3.**
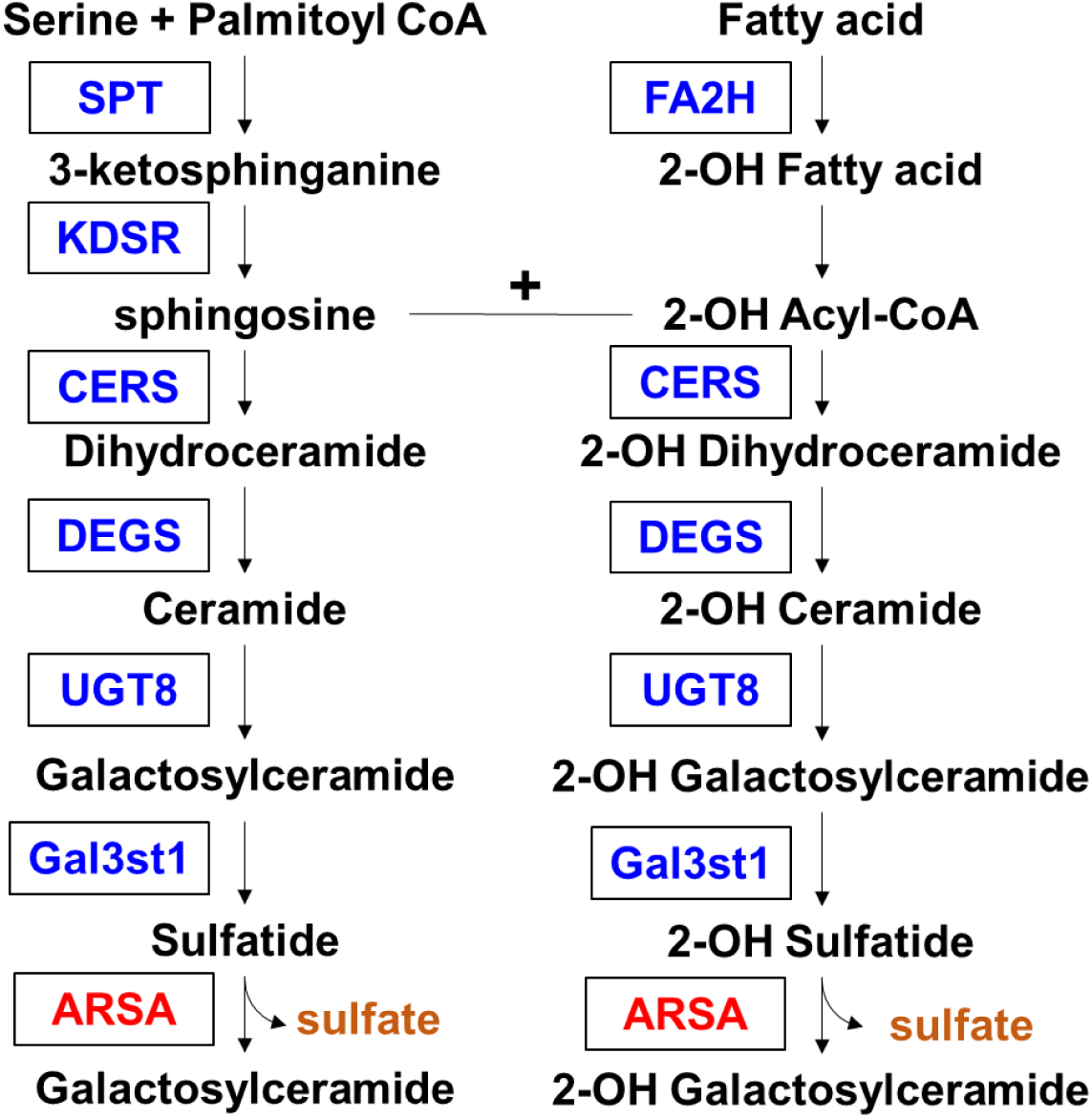
Schematic of sulfatide metabolism.

**Figure S4.**
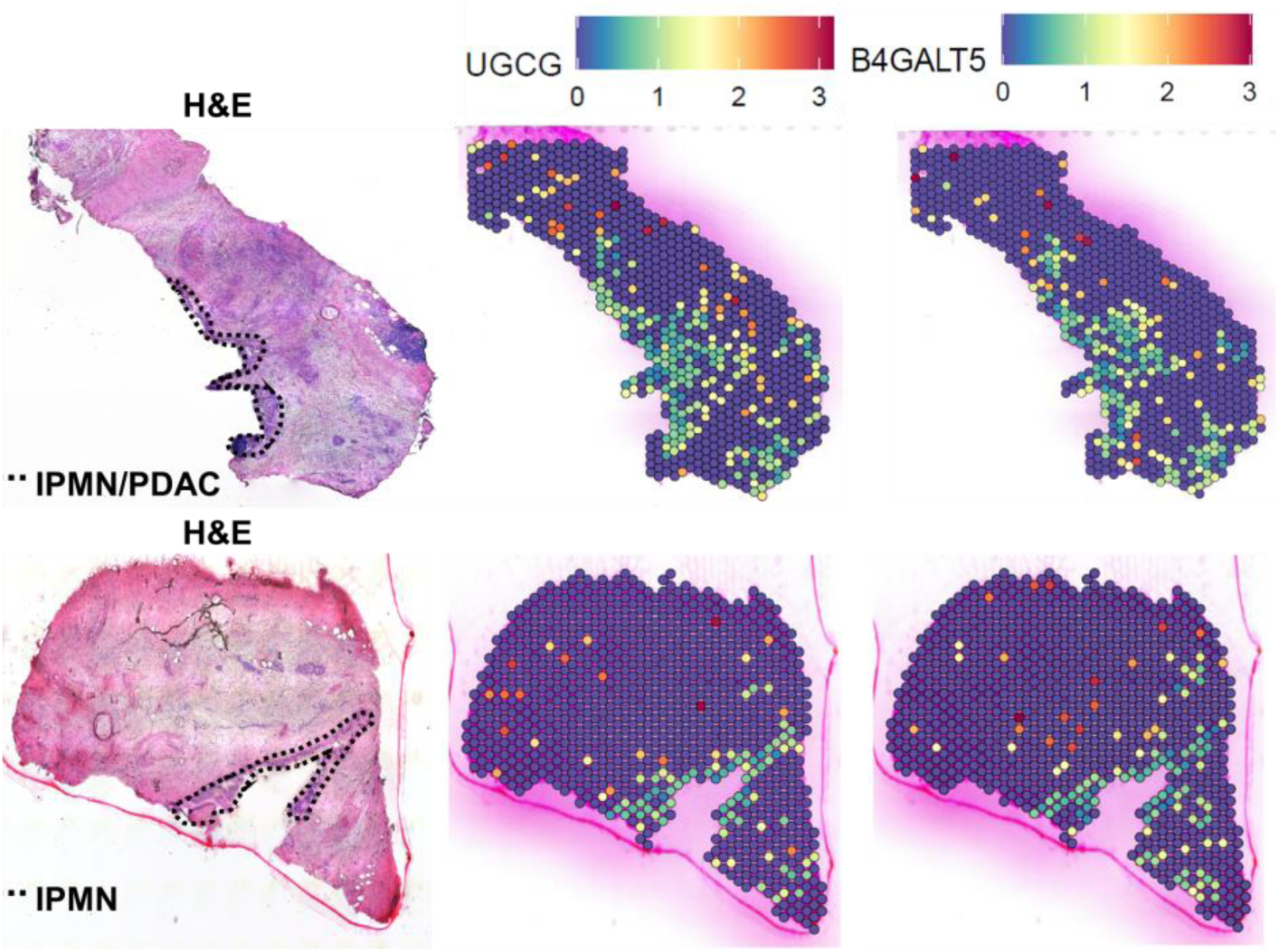
Transcript levels of glucosylceramide synthase (UGCG) and lactosylceramide synthase (encoded by B4GALT5) in IPMN tissues. The IPMN area is outlined in black.

**Figure S5.**
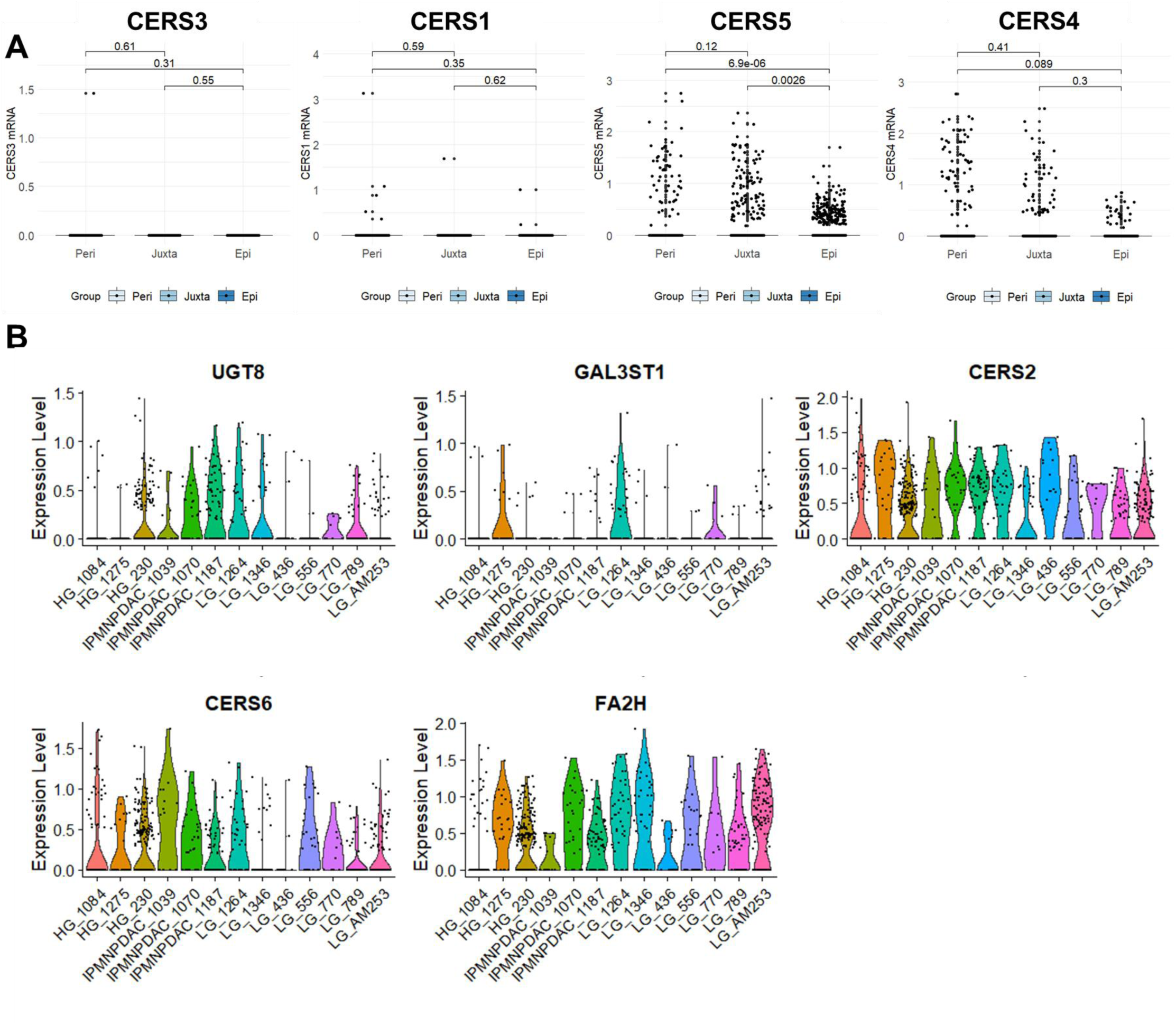
Expression levels of sulfatide-related enzymes in IPMN tissues. **A)** Boxplot showing gene expression levels for CERS1, CERS3, CERS4, and CERS5 for spots in epilesional (Epi), juxtalesional (Juxta), and perilesional (Peri) areas. Statistical significance was determined by Dunn’s multiple comparison test and adjusted P-values are reported. **B)** Violin plot showing gene expression levels for selected features for spots in epilesional areas.

**Figure S6.**
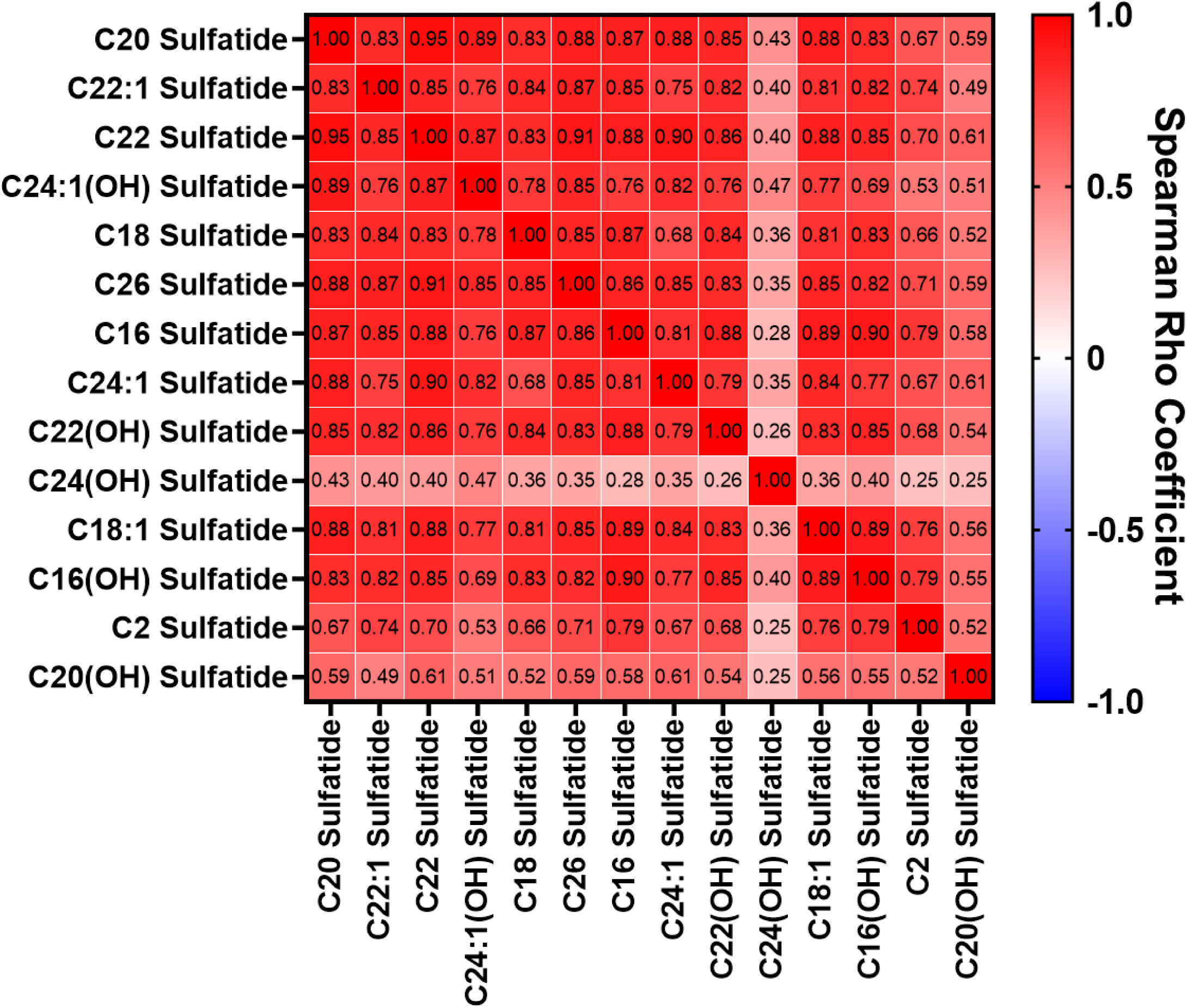
Spearman correlation matrix for sulfatides quantified in cystic fluid of patients with IPMN.

**Figure S7.**
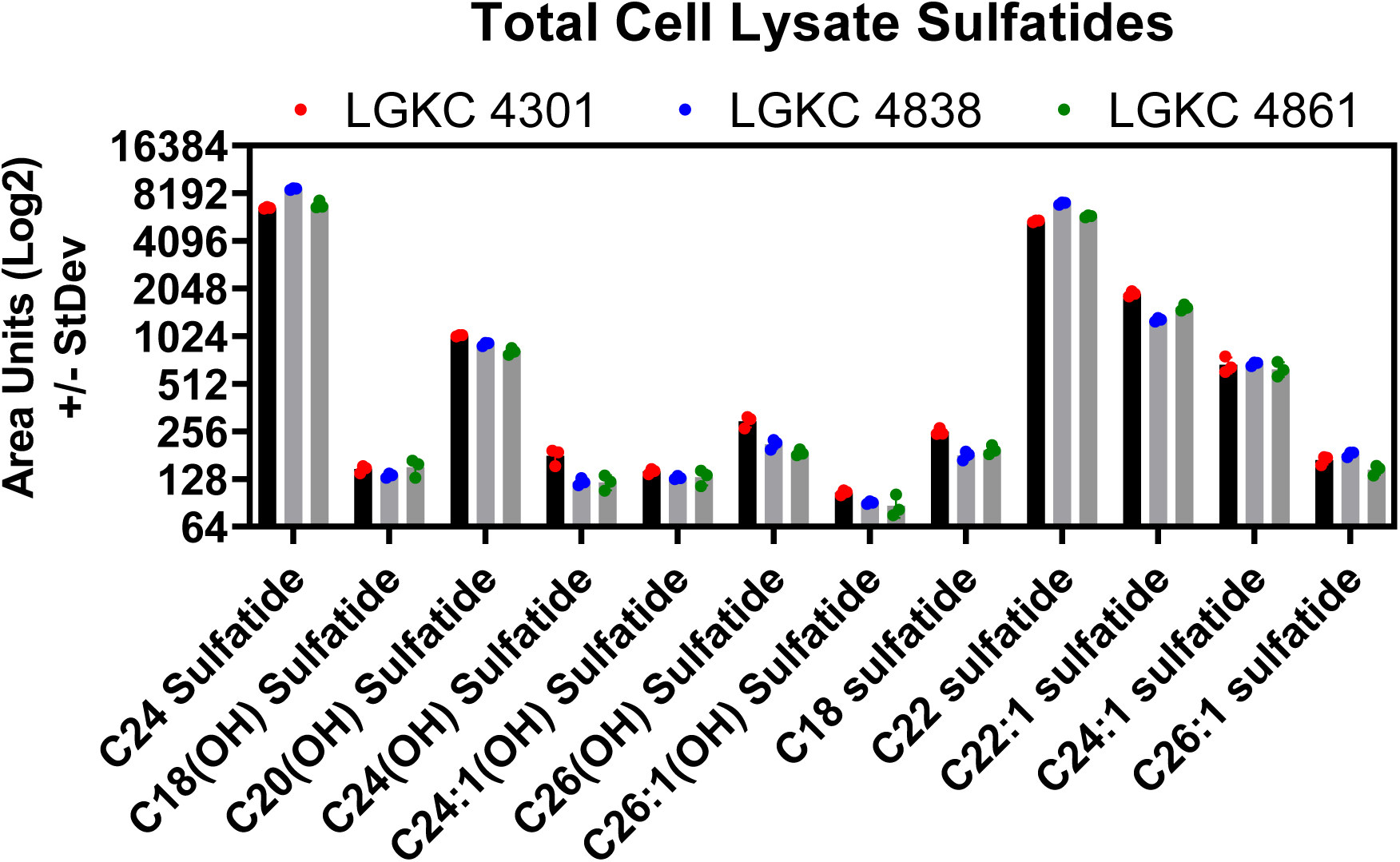
Sulfatide levels in three independent Gna^sR201C^;Kras^G12D^ activated LGKC cell lines. N =3 biological replicates were analyzed for each cell line. Values represent relative area units for each detected sulfatide species.

**Supplemental Figure S8.**
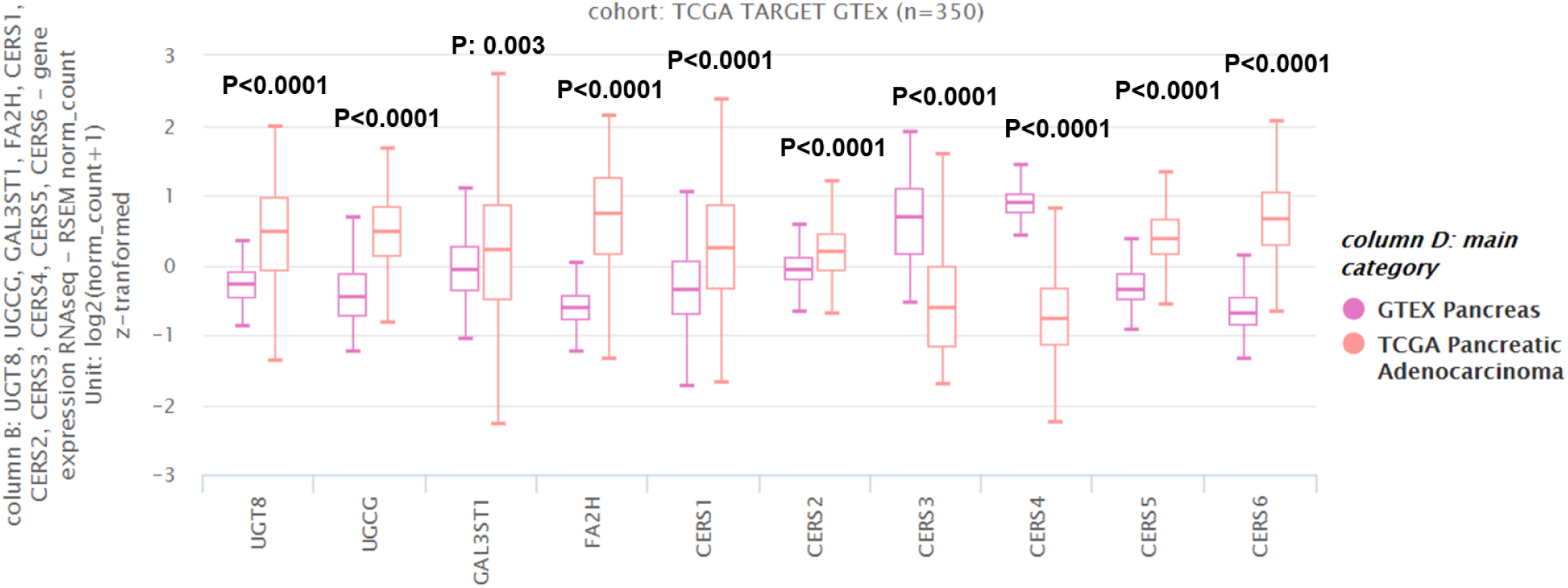
mRNA expression of sphingolipid and sulfatide related enzymes in PDAC and normal pancreas tissues. Statistical significance was determined by 2-sided Wilcoxon Rank sum test.

**Supplemental Figure S9.**
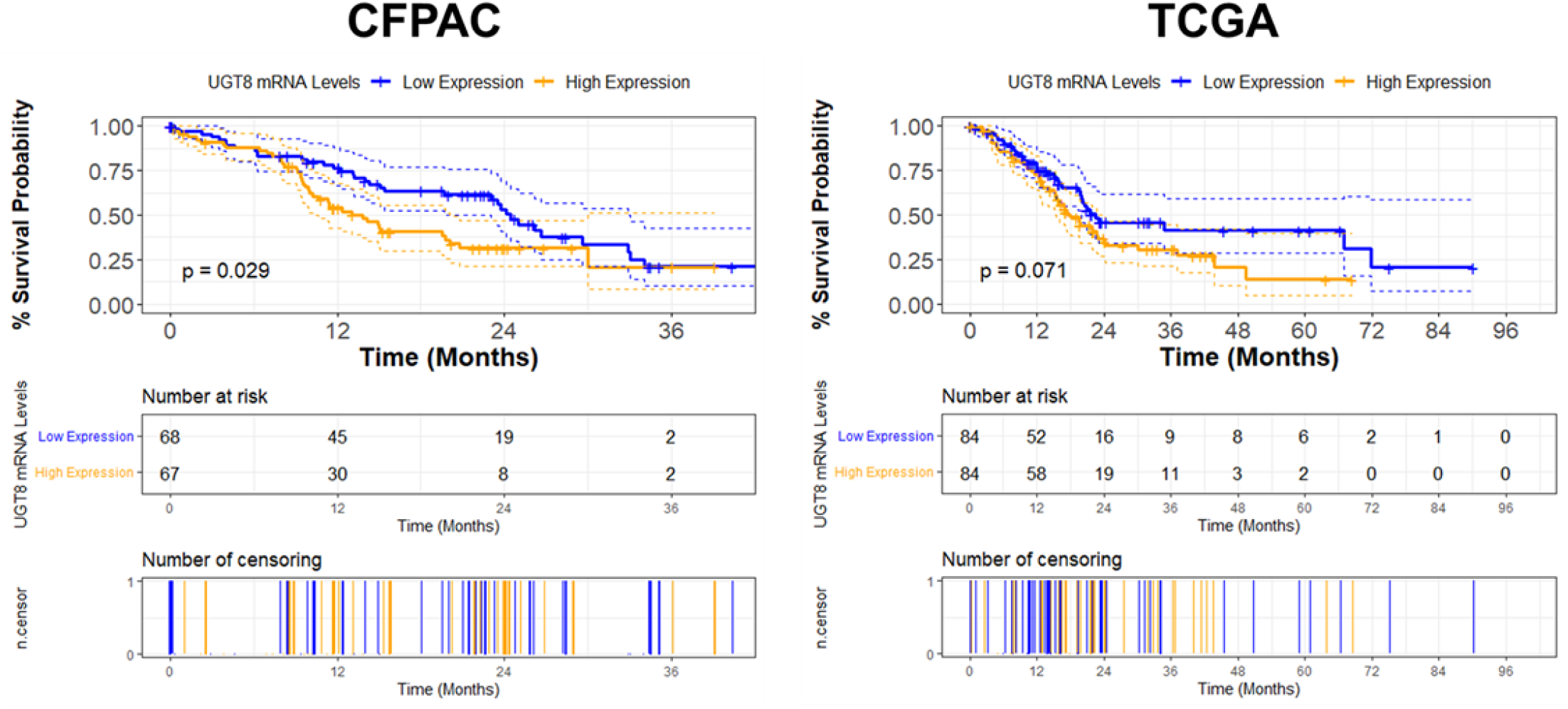
Association between tumoral UGT8 mRNA expression and overall survival in PDAC. Kaplan-Meier survival curves illustrating overall survival in PDAC cases stratified into high (> median) or low (<= median) UGT8 mRNA expression levels in the CFPAC- and TCGA-PDAC datasets. Log-rank (Mantel-Cox) tests were used to compare survival curves and 2-sided p-values are reported.

## References

1 Laffan, T. A. et al. Prevalence of unsuspected pancreatic cysts on MDCT. AJR. American journal of roentgenology 191, 802–807 (2008). https://doi.org:10.2214/ajr.07.3340

2 Megibow, A. J., Baker, M. E., Gore, R. M. & Taylor, A. The incidental pancreatic cyst. Radiologic clinics of North America 49, 349–359 (2011). https://doi.org:10.1016/j.rcl.2010.10.008

3 Salvia, R. et al. Pancreatic cystic tumours: when to resect, when to observe. European review for medical and pharmacological sciences 14, 395–406 (2010).

4 Crippa, S. et al. Mucin-producing neoplasms of the pancreas: an analysis of distinguishing clinical and epidemiologic characteristics. Clinical gastroenterology and hepatology : the official clinical practice journal of the American Gastroenterological Association 8, 213–219 (2010). https://doi.org:10.1016/j.cgh.2009.10.001

5 Noe, M. et al. Genomic characterization of malignant progression in neoplastic pancreatic cysts. Nat Commun 11, 4085 (2020). https://doi.org:10.1038/s41467-020-17917-8

6 Singhi, A. D., Koay, E. J., Chari, S. T. & Maitra, A. Early Detection of Pancreatic Cancer: Opportunities and Challenges. Gastroenterology 156, 2024–2040 (2019). https://doi.org:10.1053/j.gastro.2019.01.259

7 Lippman, S. M. et al. AACR White Paper: Shaping the Future of Cancer Prevention - A Roadmap for Advancing Science and Public Health. Cancer Prev Res (Phila) 11, 735–778 (2018). https://doi.org:10.1158/1940-6207.CAPR-18-0421

8 Singhi, A. D. & Wood, L. D. Early detection of pancreatic cancer using DNA-based molecular approaches. Nat Rev Gastroenterol Hepatol 18, 457–468 (2021). https://doi.org:10.1038/s41575-021-00470-0

9 Ying, H. et al. Oncogenic Kras maintains pancreatic tumors through regulation of anabolic glucose metabolism. Cell 149, 656–670 (2012). https://doi.org:10.1016/j.cell.2012.01.058

10 Liou, G. Y. et al. Mutant KRas-Induced Mitochondrial Oxidative Stress in Acinar Cells Upregulates EGFR Signaling to Drive Formation of Pancreatic Precancerous Lesions. Cell Rep 14, 2325–2336 (2016). https://doi.org:10.1016/j.celrep.2016.02.029

11 Carrer, A. et al. Acetyl-CoA Metabolism Supports Multistep Pancreatic Tumorigenesis. Cancer Discov 9, 416–435 (2019). https://doi.org:10.1158/2159-8290.Cd-18-0567

12 Patra, K. C. et al. Mutant GNAS drives pancreatic tumourigenesis by inducing PKA-mediated SIK suppression and reprogramming lipid metabolism. Nat Cell Biol 20, 811–822 (2018). https://doi.org:10.1038/s41556-018-0122-3

13 Encarnación-Rosado, J. & Kimmelman, A. C. Harnessing metabolic dependencies in pancreatic cancers. Nature Reviews Gastroenterology & Hepatology 18, 482–492 (2021). https://doi.org:10.1038/s41575-021-00431-7

14 Perera, R. M. & Bardeesy, N. Pancreatic Cancer Metabolism: Breaking It Down to Build It Back Up. Cancer Discov 5, 1247–1261 (2015). https://doi.org:10.1158/2159-8290.Cd-15-0671

15 Hou, P. et al. Tumor Microenvironment Remodeling Enables Bypass of Oncogenic KRAS Dependency in Pancreatic Cancer. Cancer Discov 10, 1058–1077 (2020). https://doi.org:10.1158/2159-8290.Cd-19-0597

16 Bressan, D., Battistoni, G. & Hannon, G. J. The dawn of spatial omics. Science 381, eabq4964 (2023). https://doi.org:10.1126/science.abq4964

17 Sans, M. et al. Spatial Transcriptomics of Intraductal Papillary Mucinous Neoplasms of The Pancreas Identifies NKX6-2 as a Driver of Gastric Differentiation and Indolent Biological Potential. Cancer Discov (2023). https://doi.org:10.1158/2159-8290.Cd-22-1200

18 Ideno, N. et al. GNAS(R201C) Induces Pancreatic Cystic Neoplasms in Mice That Express Activated KRAS by Inhibiting YAP1 Signaling. Gastroenterology 155, 1593–1607.e1512 (2018). https://doi.org:10.1053/j.gastro.2018.08.006

19 Sans, M. et al. Spatial Transcriptomics of Intraductal Papillary Mucinous Neoplasms of the Pancreas Identifies NKX6-2 as a Driver of Gastric Differentiation and Indolent Biological Potential. Cancer Discovery 13, 1844–1861 (2023). https://doi.org:10.1158/2159-8290.Cd-22-1200

20 Fahrmann, J. F. et al. A Plasma-Derived Protein-Metabolite Multiplexed Panel for Early-Stage Pancreatic Cancer. J Natl Cancer Inst 111, 372–379 (2019). https://doi.org:10.1093/jnci/djy126

21 Fahrmann, J. F. et al. A MYC-Driven Plasma Polyamine Signature for Early Detection of Ovarian Cancer. Cancers (Basel) 13 (2021). https://doi.org:10.3390/cancers13040913

22 Fahrmann, J. F. et al. Association Between Plasma Diacetylspermine and Tumor Spermine Synthase With Outcome in Triple-Negative Breast Cancer. J Natl Cancer Inst 112, 607–616 (2020). https://doi.org:10.1093/jnci/djz182

23 Vykoukal, J. et al. Caveolin-1-mediated sphingolipid oncometabolism underlies a metabolic vulnerability of prostate cancer. Nat Commun 11, 4279 (2020). https://doi.org:10.1038/s41467-020-17645-z

24 Cao, L. et al. Proteogenomic characterization of pancreatic ductal adenocarcinoma. Cell 184, 5031–5052.e5026 (2021). https://doi.org:10.1016/j.cell.2021.08.023

25 Gao, J. et al. Integrative analysis of complex cancer genomics and clinical profiles using the cBioPortal. Sci Signal 6, pl1 (2013). https://doi.org:10.1126/scisignal.2004088

26 Chen, K. et al. Immune profiling and prognostic model of pancreatic cancer using quantitative pathology and single-cell RNA sequencing. J Transl Med 21, 210 (2023). https://doi.org:10.1186/s12967-023-04051-4

27 Hao, Y. et al. Integrated analysis of multimodal single-cell data. Cell 184, 3573–3587 e3529 (2021). https://doi.org:10.1016/j.cell.2021.04.048

28 Grambsch, P. & Therneau, T. Proportional Hazards Tests and Diagnostics Based on Weighted Residuals Biometrika, 81, 515–526. Find this article online (1994).

29 Bosio, A., Binczek, E., Le Beau, M. M., Fernald, A. A. & Stoffel, W. The human gene CGT encoding the UDP-galactose ceramide galactosyl transferase (cerebroside synthase): cloning, characterization, and assignment to human chromosome 4, band q26. Genomics 34, 69–75 (1996). https://doi.org:10.1006/geno.1996.0242

30 Blomqvist, M., Zetterberg, H., Blennow, K. & Månsson, J.-E. Sulfatide in health and disease. The evaluation of sulfatide in cerebrospinal fluid as a possible biomarker for neurodegeneration. Molecular and Cellular Neuroscience 116, 103670 (2021). https://doi.org:https://doi.org/10.1016/j.mcn.2021.103670

31 Iglesias-Bartolome, R. et al. Inactivation of a Galpha(s)-PKA tumour suppressor pathway in skin stem cells initiates basal-cell carcinogenesis. Nature cell biology 17, 793–803 (2015). https://doi.org:10.1038/ncb3164

32 Sentelle, R. D. et al. Ceramide targets autophagosomes to mitochondria and induces lethal mitophagy. Nat Chem Biol 8, 831–838 (2012). https://doi.org:10.1038/nchembio.1059

33 Cao, Q. et al. Inhibition of UGT8 suppresses basal-like breast cancer progression by attenuating sulfatide-αVβ5 axis. J Exp Med 215, 1679–1692 (2018). https://doi.org:10.1084/jem.20172048

34 Yoda, Y. et al. Glycolipids in human lung carcinoma of histologically different types. J Natl Cancer Inst 63, 1153–1160 (1979).

35 Morichika, H., Hamanaka, Y., Tai, T. & Ishizuka, I. Sulfatides as a predictive factor of lymph node metastasis in patients with colorectal adenocarcinoma. Cancer 78, 43–47 (1996). https://doi.org:10.1002/(sici)1097-0142(19960701)78:1<43::Aid-cncr8>3.0.Co;2-i

36 Makhlouf, A. M., Fathalla, M. M., Zakhary, M. A. & Makarem, M. H. Sulfatides in ovarian tumors: clinicopathological correlates. Int J Gynecol Cancer 14, 89–93 (2004). https://doi.org:10.1111/j.1048-891x.2004.014223.x-1

37 Garcia, J., Callewaert, N. & Borsig, L. P-selectin mediates metastatic progression through binding to sulfatides on tumor cells. Glycobiology 17, 185–196 (2007). https://doi.org:10.1093/glycob/cwl059

38 Duchesneau, P., Gallagher, E., Walcheck, B. & Waddell, T. K. Up-regulation of leukocyte CXCR4 expression by sulfatide: an L-selectin-dependent pathway on CD4+ T cells. Eur J Immunol 37, 2949–2960 (2007). https://doi.org:10.1002/eji.200737118

39 Sandhoff, R. et al. Chemokines bind to sulfatides as revealed by surface plasmon resonance. Biochim Biophys Acta 1687, 52–63 (2005). https://doi.org:10.1016/j.bbalip.2004.11.011

40 Popovic, Z. V. et al. Sulfated glycosphingolipid as mediator of phagocytosis: SM4s enhances apoptotic cell clearance and modulates macrophage activity. J Immunol 179, 6770–6782 (2007). https://doi.org:10.4049/jimmunol.179.10.6770

41 Altomare, E. et al. Synthetic isoforms of endogenous sulfatides differently modulate indoleamine 2,3-dioxygenase in antigen presenting cells. Life Sci 89, 176–181 (2011). https://doi.org:10.1016/j.lfs.2011.05.015

42 Amobi, A., Qian, F., Lugade, A. A. & Odunsi, K. Tryptophan Catabolism and Cancer Immunotherapy Targeting IDO Mediated Immune Suppression. Adv Exp Med Biol 1036, 129–144 (2017). https://doi.org:10.1007/978-3-319-67577-0_9

43 Platten, M., Nollen, E. A. A., Röhrig, U. F., Fallarino, F. & Opitz, C. A. Tryptophan metabolism as a common therapeutic target in cancer, neurodegeneration and beyond. Nat Rev Drug Discov 18, 379–401 (2019). https://doi.org:10.1038/s41573-019-0016-5

44 Lemay, A. M. et al. High FA2H and UGT8 transcript levels predict hydroxylated hexosylceramide accumulation in lung adenocarcinoma. J Lipid Res 60, 1776–1786 (2019). https://doi.org:10.1194/jlr.M093955

45 Zhou, X. et al. Dysregulated ceramides metabolism by fatty acid 2-hydroxylase exposes a metabolic vulnerability to target cancer metastasis. Signal Transduction and Targeted Therapy 7, 370 (2022). https://doi.org:10.1038/s41392-022-01199-1

